# *In Vitro* Dynamic and Quantitative Monitoring of Strigolactone-signaling Complex Formation by Time-resolved FRET

**DOI:** 10.1101/2025.11.18.688998

**Authors:** Taiki Suzuki, Kotaro Nishiyama, Yusuke Kato, Chihiro Shinkai, Tomoya Ishikawa, Jekson Robertlee, Michio Kuruma, Shinya Hagihara, Marco Bürger, Kosuke Fukui, Tadao Asami, Yoshiya Seto

## Abstract

Strigolactones (SLs) are a class of plant hormones that play a critical role in the suppression of shoot branching. Furthermore, they are exuded from roots and act as signaling molecules for inter-organism communication in the rhizosphere. Strigolactones trigger those responses by inducing protein–protein interactions (PPIs) of signaling components and subsequent proteolysis of transcriptional repressors. The sequential event involves SL hydrolysis mediated by SL receptors belonging to an α/μ-hydrolase family, although the physiological role of SL hydrolysis is a subject of debate. To date, SL-induced PPIs have been analyzed by methods such as yeast-two hybrid, pull-down, and AlphaScreen assays. However, the kinetic aspect of PPI profiles has not been well studied. Here, we developed an *in vitro* method to monitor the formation of the SL signaling complex based on Time-Resolved Förster Resonance Energy Transfer (TR-FRET) technology. Our TR-FRET-based assay system allows us to analyze the mode of action of SL analogs from kinetic and quantitative perspectives. Notably, our method revealed differences in the intensity and time-dependency of PPI signals among different SL analogs with a range of hydrolyzabilities. In addition, we found that tolfenamic acid, an antagonist of the SL receptor, inhibited the SL-induced PPI but could not disrupt the already-formed signaling complex. The TR-FRET system was also used to rapidly and specifically detect naturally occurring SLs from root exudates containing many impurities. This work provides insights into the molecular mechanism of SL perception as well as a powerful tool for activity-based screening of SL signaling modulators.

**Significance statement:** The dual roles of strigolactone (SL) receptors in both the perception and deactivation of SLs make it difficult to elucidate the underlying molecular mechanism of SL signaling. We developed a new *in vitro* method to evaluate the dynamic activation of the SL receptor, and used it to gain deeper insights into the molecular mechanism of SL-signaling complex formation in response to the SL receptor agonists.

## Introduction

Plant hormones regulate every aspect of plant life including growth, development, and responses to environmental stresses (Santner *et al*., 2009). Most plant hormones promote protein–protein interactions (PPIs) between receptors and signaling components (Takeuchi *et al*., 2021). The PPIs induce each downstream signal through protein phosphorylation or degradation. Thus, the development of assay methods to monitor hormone-induced PPIs, the key step in plant hormone signaling, is critical for thoroughly understanding the hormone signaling mechanism.

Strigolactones (SLs) are the most recently identified class of plant hormones. They play an important role in the inhibition of shoot branching in angiosperms (Umehara *et al*., 2008; Gomez-Roldan *et al*., 2008). In *Arabidopsis thaliana*, the SL receptor AtDWARF14 (AtD14) recognizes SL and subsequently recruits the transcriptional repressor proteins SUPPRESSOR OF MAX2 1-LIKE 6, 7, and 8 (SMXL6/7/8) and the F-box protein MORE AXIRALLY GROWTH 2 (MAX2), which is a component of the SCF E3 ubiquitin ligase complex, *via* physical interactions. After the ubiquitination of the repressor proteins, they are degraded through the 26S proteasome system, leading to the inhibition of shoot branching (Wang *et al*., 2015; Soundappan *et al*., 2015; Yao *et al*., 2016; Zhao *et al*., 2015; Zhou *et al*., 2013; Jiang *et al*., 2013). Various SLs also play important roles in the rhizosphere as symbiotic signals for arbuscular mycorrhizal fungi (Akiyama *et al*., 2005). The SLs exuded from roots are exploited as germination stimulants by obligate root parasitic plants in Orobanchaceae family, which pose a serious threat to global food security (Cook *et al*., 1966). Proteins in the divergent clade KARRIKIN INSENSITIVE2/HYPOSENSITIVE TO LIGHT (KAI2/HTL), which is a paralogous family of AtD14, hereinafter called KAI2d, were identified as the receptors for host root-derived SLs in the obligate root parasitic plant *Striga hermonthica* (Tsuchiya *et al*., 2015; Toh *et al*., 2015; Conn *et al*., 2015). More recently, highly sensitive SL receptors were isolated and characterized from another obligate parasitic plant, *Orobanche minor*, and from the facultative hemiparasitic plant, *Phtheirospermum japonicum* (Takei *et al*., 2023; Takei *et al*., 2024). Similar to AtD14, SL-dependent activation of KAI2d promotes PPIs among KAI2d, SCF^MAX2^, and the transcriptional repressor protein SUPPRESSOR OF MAX2 1 (SMAX1), leading to the degradation of SMAX1 through the ubiquitin–proteasome system (Liu *et al*., 2014; Stanga *et al*., 2013; Soundappan *et al*., 2015; Yao *et al*., 2017; Nelson, 2021).

Strigolactone receptors, which belong to the α/μ-hydrolase superfamily, not only interact with SL, but also hydrolyze it (Hamiaux *et al*., 2012; Seto *et al*., 2019). Recent biochemical and structural studies revealed that the SL hydrolysis intermediate covalently binds to AtD14 catalytic residues in the signaling complex, indicating that SL hydrolysis is indispensable for the activation of AtD14 (Yao *et al*., 2016; Bürger and Chory, 2020). However, the AtD14^D218A^ mutant, which has no SL hydrolysis activity because it lacks the catalytic triad, retained the ability to interact with AtSMXL7 in a SL-dependent manner and restored the *atd14* mutant phenotype (Seto *et al*., 2019). Additionally, studies using chemical and biological methods indicated that the full activation of SL receptors depends on the hydrolysis of SL agonists, but the hydrolysis process is not necessarily required for SL signal transduction (Wang *et al*., 2022; Uraguchi *et al*., 2018; Yasui *et al*., 2019). Despite these advances, the exact function of SL hydrolysis and the timing of the formation of the complex are not completely understood. Notably, an analysis of the dynamics of SL-dependent complex formation associated with the hydrolysis process would provide new insights into the SL signaling mechanism. So far, the PPIs of SL signaling components have been detected using yeast-two hybrid, pull-down, and AlphaScreen assays. However, these methods are not suitable for kinetic analyses (Zhao *et al*., 2015; Wang *et al*., 2021; Hamiaux *et al*., 2012; Zhou *et al*., 2013).

Here, we developed the Time-Resolved Förster Resonance Energy Transfer (TR-FRET) assay system to quantitatively and kinetically detect the formation of the SL signaling complexes derived from *A. thaliana* and *O. minor*. The TR-FRET signal arises from the proximity between a luminescent lanthanide complex and a typical fluorescent dye (e.g. fluorescein, Cy5, or GFP). The physical properties of the lanthanide complexes, which have long lifetimes, sharp emission spectra, and large Stokes shift, enable the quantitative and kinetic analysis of PPIs, as well as the characterization of agonist/antagonist activities. Using the TR-FRET assay system, we were able to analyze the mode of action of SL receptor ligands. In addition, this method revealed that the SL receptor–repressor complex is so kinetically stable that an antagonist could not disrupt the PPIs within the already-formed complex. Time-course PPI analyses exhibited variations in intensity and time-dependency of the PPI signals among various SL analogs with different degrees of resistance to receptor-mediated hydrolysis. Overall, our results show that this novel technique can provide in-depth insights into the molecular mechanisms of the formation of the SL receptor complex. In addition to its use in mechanism-of-action studies, the TR-FRET assay system can rapidly and specifically detect naturally occurring SLs in root exudate mixtures. This selective activity-based probing method will facilitate the discovery of natural SL receptor ligands that control the signaling pathway.

## Results

### Development of the TR-FRET assay system to detect SL signaling complex formation

To monitor the PPIs of SL signaling components, we set out to develop the TR-FRET assay system using the protein pairs AtD14–AtMAX2 and AtD14–AtSMXL7. We chose N- or C-terminal mEGFP-fused AtD14 with a 6×HN tag (mEGFP-AtD14 or AtD14-mEGFP, respectively) as the acceptor for the TR-FRET assay. The N-terminal 6×His-3×FLAG-tagged AtMAX2 was transiently expressed in *Nicotiana benthamiana* and purified by Co^2**+**^-affinity chromatography (Figure S1A,B). Based on a previous report, we expressed N-terminal 3×FLAG- and C-terminal 6×His-tagged AtSMXL7 in *Escherichia coli*, targeted to the periplasmic fraction using the pelB signal sequence (Figure S1C,D) (Khosla *et al*., 2020). Although the purified samples contained some impurities as well as AtMAX2/AtSMXL7, the pull-down assays showed that AtD14–AtMAX2 and AtD14–AtSMXL7 interactions proceeded in the presence of *racemic* GR24 (*rac*-GR24, a mixture of (+)-GR24 and (−)-GR24), a synthetic agonist of the SL receptor (Figure 1A). These results confirm that the prepared proteins retained their original activities.

**Figure 1.**
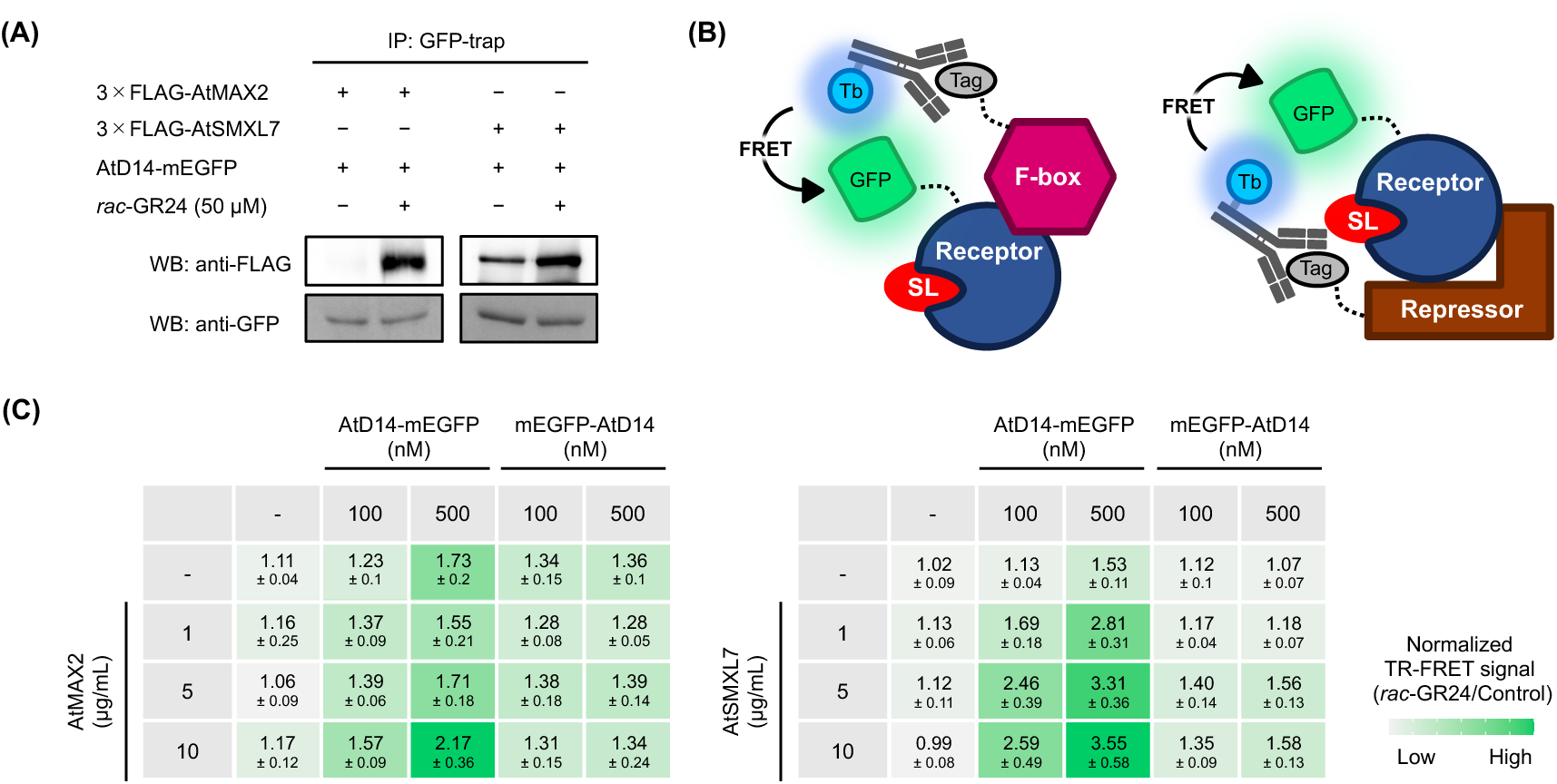
(A) Pull-down assay to detect SL-induced complex formation of AtD14– AtMAX2 and AtD14–AtSMXL7 pairs. AtD14-mEGFP was detected by anti-GFP antibody. AtMAX2 and AtSMXL7 were detected by anti-FLAG antibody. IP, immunoprecipitation. WB, western blotting. (B) Schematic of TR-FRET assay for measuring SL-induced PPIs. The receptor protein is fused with mEGFP (GFP). The F-box and repressor proteins are FLAG-tagged and labeled by anti-FLAG antibody harnessing terbium-cryptate (Tb). SL induces each signaling complex resulting in a TR-FRET signal enhancing. (C) The dependency of each SL-signaling component in AtD14–AtMAX2 and AtD14–AtSMXL7 TR-FRET assays. Heatmap-projected values represent fold changes of TR-FRET signal upon treatment with *rac*-GR24 (50 µM) at a 300 min incubation period. Data are the means ± SD (*n* = 3). Values of TR-FRET signals before normalization are shown in Figure S2A,B.

Next, we performed TR-FRET assays using the same proteins at a range of concentrations. In these assays, we conducted *in situ* labeling of 3×FLAG-tagged AtMAX2 or AtSMXL7 with anti-FLAG antibodies connecting the terbium-cryptate FRET donors (Figure 1B). We measured TR-FRET signals, that indicate the ratio of the fluorescence intensity of the donor Tb-cryptate (485 nm) to that of the acceptor mEGFP (520 nm) in the absence or presence of *rac*-GR24 (Figure S2A,B). Figure 1C displayed TR-FRET signals upon addition of *rac*-GR24 normalized by the buffer control. The *rac*-GR24-dependent signals clearly increased in the presence of almost all pairs of proteins (Figure 1C). We found that AtD14-mEGFP yielded higher SL-dependent TR-FRET signals than mEGFP-AtD14 with both partner proteins (Figure 1C). In the presence of AtD14-mEGFP (100 nM), the SL-dependent TR-FRET signal slightly increased from 1.23 to 1.57 upon addition of AtMAX2 (Figures 1C). In the case of the AtD14–AtSMXL7 pair, the addition of AtSMXL7 raised the SL-dependent signal from 1.13 to 2.59 in the presence of 100 nM of AtD14-mEGFP (Figure 1C). In contrast, partner protein-independent nonspecific signals were observed in the presence of 500 nM AtD14. In the following experiments, therefore, we used AtD14-mEGFP (100 nM) as the acceptor because sufficient SL-triggered PPI signals were detected with this concentration. To confirm that the signal increase in the presence of *rac*-GR24 was derived from the structural activation of AtD14, TR-FRET assays were performed by using the catalytic triad mutant, AtD14^S97A^, which has been reported to lose both the catalytic and signaling function (Hamiaux *et al*., 2012; Seto *et al*., 2019). No signal induction was observed in either AtD14^S97A^–AtMAX2 or the AtD14^S97A^–AtSMXL7 upon addition of enough *rac*-GR24 (Figure S3A,B). To further examine the specificity of these signals, we prepared 6×His-AtMAX2 and StrepII-HiBiT-AtSMXL7-6×His without the anti-FLAG epitope tags (Figure S3C,D,E,F). As expected, the SL-dependent PPI signals were not increased in a protein-dependent manner, even in the presence of Tb cryptate-modified anti-FLAG antibodies (Figure S3G,I). In addition, these negative control proteins competed with the 3×FLAG-tagged AtMAX2 and AtSMXL7 (Figure S3H,J). These results show that our TR-FRET assay system specifically detects SL-dependent PPI behavior in AtD14–AtMAX2 and AtD14–AtSMXL7. We used the AtD14–AtSMXL7 pair in the following experiments because of its higher signal to noise ratio compared with that of AtD14–AtMAX2.

### Quantitative and kinetic analysis of complex formation by the TR-FRET assay system

It has been reported that (+)-GR24, the 2*’R* stereoisomer of GR24, more strongly inhibits shoot branching in SL biosynthesis mutants of Arabidopsis than does (−)-GR24, the 2*’S* stereoisomer (Scaffidi *et al*., 2014). Despite the distinct biological activities of the enantiomer pair, in *in vitro* analyses, namely, differential scanning fluorometry (DSF), there were no significant differences in the thermal stability shift in the presence of each stereoisomer (Figure S4), consistent with the results of a previous study (Scaffidi *et al*., 2014; Waters *et al*., 2015). To investigate the difference in the mode of action between (+)-GR24 and (−)-GR24, quantitative and kinetic TR-FRET assays using the AtD14–AtSMXL7 pair were carried out. Time-course measurements showed gradual increases in PPI signals in both cases (Figure 2A). However, the maximum intensity of the PPI signal was significantly different between (+)-GR24 and (−)-GR24, in spite of their comparable 50% effective concentration (EC50) values (0.30 and 0.68 µM, respectively) (Figure 2B). These results show that (−)-GR24 hardly activates the PPI, although it is recognized by the receptor.

**Figure 2.**
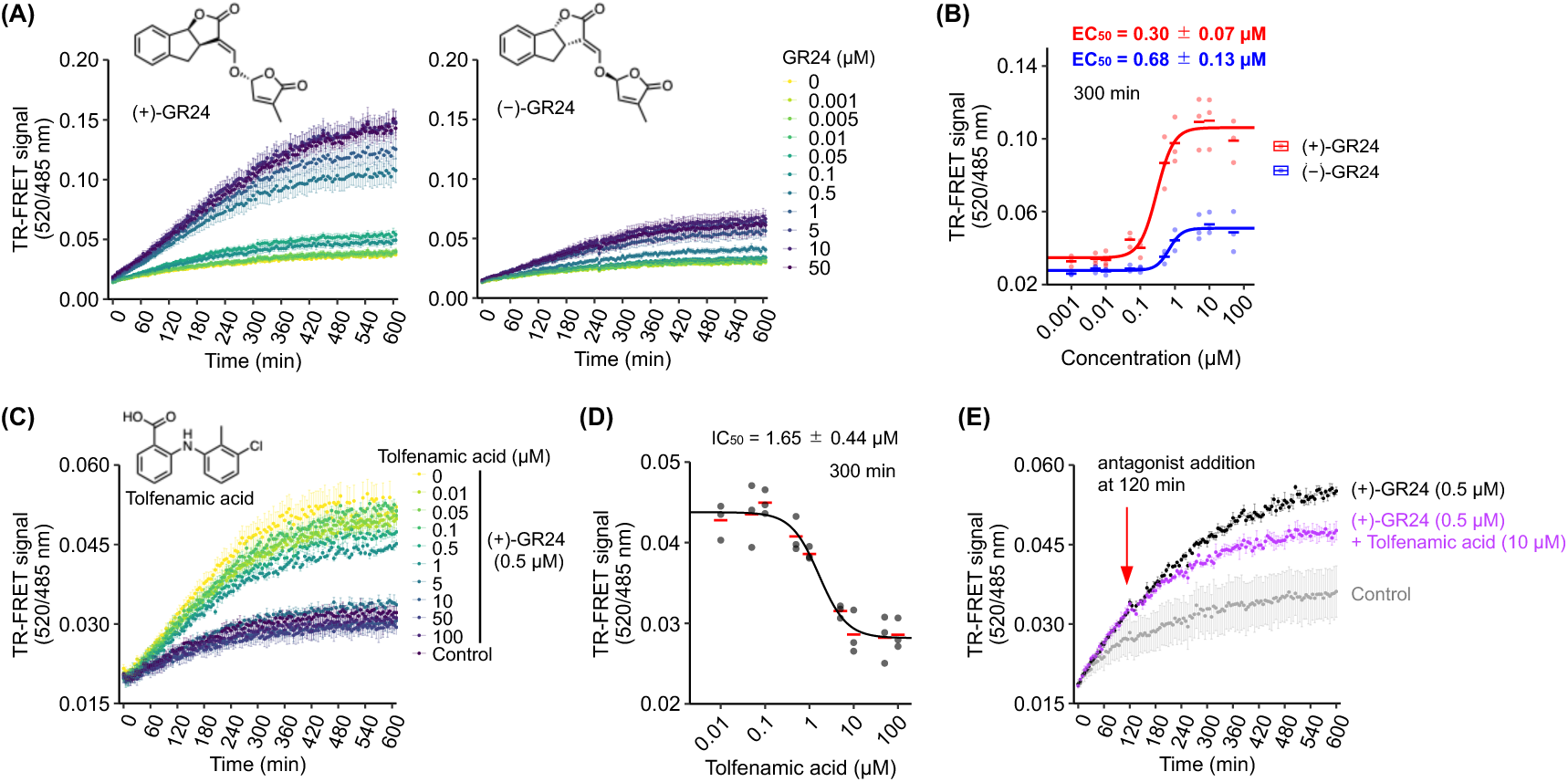
Quantitative and kinetic analysis of the AtD14 agonist and antagonist activities in TR-FRET assays. (A) Kinetic measurements of AtD14–SMXL7 TR-FRET signals in the presence of indicated concentration of (+)-GR24 and (−)-GR24. The assays were performed using AtD14-mEGFP (100 nM) and 3×FLAG-AtSMXL7 (5 µg/mL). Data are the means ± SE (*n* = 3). (B) Dose-titration of (+)-GR24 and (−)-GR24 in AtD14–AtSMXL7 TR-FRET assays at the single time point (300 min) extracted from the data as shown in Figure 2A. Dots represent individual data values of three technical replicates. (C) Kinetic analysis of antagonistic activity of tolfenamic acid on GR24-induced AtD14–AtSMXL7 complex formation. The assay was performed using AtD14-mEGFP (100 nM) and 3×FLAG-AtSMXL7 (5 µg/mL). Control was performed without (+)-GR24 and tolfenamic acid. Data are the means ± SE (*n* = 3). (D) Dose-response curve of tolfenamic acid in TR-FRET inhibition assay with GR24-induced AtD14–AtSMXL7 at the single time point (300 min) extracted from the data as shown in Figure 2C. Dots represent individual data values of three technical replicates. (E) The antagonistic effect of tolfenamic acid after the formation of AtD14–AtSMXL7 complex. The red arrow indicates the timing of tolfenamic acid treatment. The assay was performed using AtD14-mEGFP (100 nM) and 3×FLAG-AtSMXL7 (5 µg/mL). Data are the means ± SE (*n* = 3).

Next, to evaluate the applicability of the TR-FRET assay system, we investigated the antagonistic activity of tolfenamic acid, which is an antagonist of the SL receptors of petunia, rice, and Arabidopsis (Hamiaux *et al*., 2018). When tolfenamic acid was supplied alongside (+)-GR24 to the AtD14–AtSMXL7 pair, the increase in the TR-FRET signal was clearly inhibited (Figure 2C). As shown in Figure 2D, the 50% inhibitory concentration (IC50) value at 300 min was calculated as 1.65 µM, similar to the *Ki* value against AtD14 (2.52 µM) (Hamiaux *et al*., 2018). To further examine whether this SL receptor antagonist could disrupt the already-formed SL signaling complex, tolfenamic acid was added after the protein pair was pre-incubated with (+)-GR24 for 120 min. Surprisingly, while the increase in the TR-FRET signal stopped immediately upon addition of the antagonist, the elevated signal barely decreased (Figure 2E). These results suggest that the AtD14–AtSMXL7 complex is kinetically stable and/or the antagonist cannot access the catalytic pocket of the receptor in the ternary complex.

### Adaptation of TR-FRET assay system to *O. minor* SL signaling components

Next, to expand the applicability of this system, we developed the TR-FRET assay system for the *O. minor* receptor–repressor protein pair. Another study conducted a functional analysis of SL receptors in *O. minor*, and found that OmKAI2d3, one of the *KAI2d* gene products, was highly sensitive to SL (Takei *et al*., 2023). We considered that quantitative and kinetic TR-FRET analyses would provide new insights into this sensitivity. Because the SMAX1 homolog in *O. minor* has not been identified, we searched for a SMAX1-like gene (KAL6540832.1) in the entire *O. minor* genome (Bürger *et al*., 2025). As shown in the phylogenetic tree, the amino acid sequence of this gene product had high homology with SMAX1 and SMXL2 from *A. thaliana* (Figure S5). Thus, we named this gene *OmSMAX1* and cloned its coding sequence from the cDNA of *O. minor*. Initially, 6×HN-tagged OmKAI2d3-mEGFP and 3×FLAG-OmSMAX1-6×His were prepared in the same way as the corresponding proteins of *A. thaliana,* but almost all of the full-length OmSMAX1 protein was degraded after affinity column purification (Figure S6A,B). Because the previous study showed that the D1M-domain of SMAX1 is an important region for ligand-induced PPI, we prepared the D1M domain of OmSMAX1 (OmSMAX1-D1M) as an alternative to the full-length protein to improve its stability (Figure S6C,D) (Khosla *et al*., 2020). The results of a pull-down assay showed that the PPI between OmKAI2d3 and OmSMAX1-D1M was enhanced by *rac*-GR24, indicating that the D1M domain is also a critical region for the SL-dependent PPI between OmKAI2d3 and OmSMAX1 (Figure 3A). Next, we performed the TR-FRET assay using these proteins and found that SL-dependent TR-FRET signals significantly increased in OmKAI2d3- and OmSMAX1-D1M-dependent manners (Figure 3B, Figure S7). In the presence of 100 nM OmKAI2d3-mEGFP and 5 µg/mL 3×FLAG-OmSMAX1-D1M, the TR-FRET signal increased 2.36-fold upon *rac*-GR24 treatment, with a low OmSMAX1-D1M-independent signal. Accordingly, we used these conditions for subsequent experiments.

**Figure 3.**
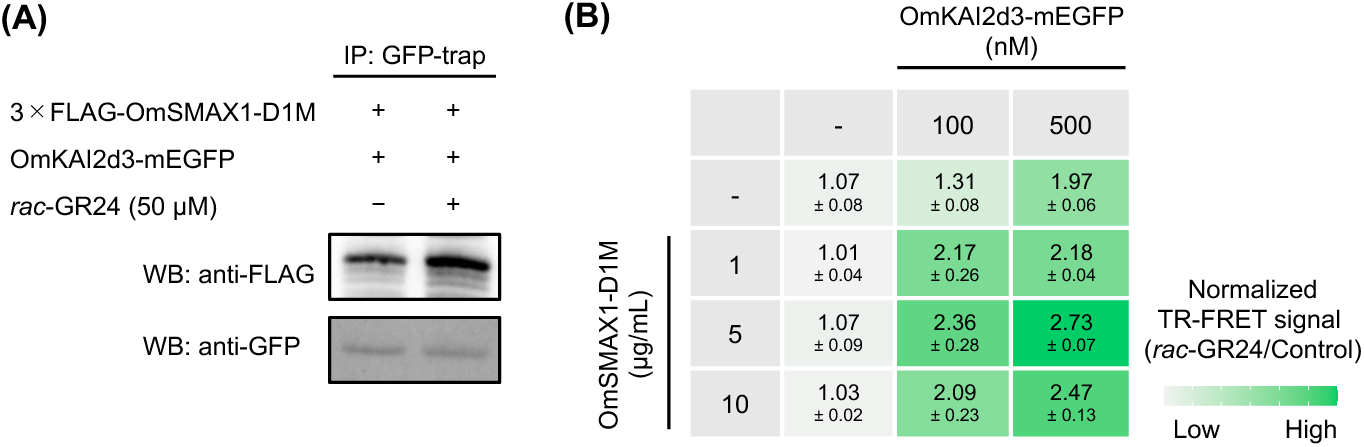
Adaptation of the TR-FRET assay system to the homologs’ pair of O. minor. (A) Pull-down assay to detect SL-induced complex formation between OmKAI2d3 and OmSMAX1 D1M domain. OmKAI2d3-mEGFP was detected by anti-GFP antibody. OmSMAX1-D1M was detected by anti-FLAG antibody. IP, immunoprecipitation. WB, western blotting. (B) The dependency of each SL-signaling component in TR-FRET assays. Heatmap-projected values represent fold changes of TR-FRET signal upon treatment with *rac*-GR24 (50 µM) at a 300 min incubation period. Data are the means ± SD (*n* = 3). Values of TR-FRET signals before normalization are shown in Figure S7. The data in the absence of OmSMAX1 are identical to those in Figure 9S.

To further validate the TR-FRET profiles, we performed the assays using (+)-GR24 and (−)-GR24. The results showed that (+)-GR24 more strongly increased the PPI signal of OmKAI2d3–OmSMAX1-D1M than did (−)-GR24 in a time-dependent manner (Figure 4A). As measured at a single time point, the EC50 value of (+)-GR24 for complex formation was 0.046 µM, and that of (−)-GR24 was about 1000-times higher (Figure 4B). Previous studies showed that the 2’*R* configuration at the methyl butenolide ring is important for the agonistic activity of SL analogs on germination of root parasitic plants (Umehara *et al*., 2015; Scaffidi *et al*., 2014). Consistent with the results of previous studies, our *in vivo* germination assays indicated that our *O. minor* seeds responded more strongly to (+)-GR24 than to (−)-GR24 (Figure S8A). To analyze the direct interaction between these chemicals and OmKAI2d3, we performed a DSF assay. The results showed that (+)-GR24 significantly decreased the melting temperature of OmKAI2d3, but (−)-GR24 did not (Figure S8B). These results demonstrate that our TR-FRET assay system can specifically and quantitatively evaluate the activity of SL agonists to induce PPIs among the SL signaling components of *O. minor,* just like in *A. thaliana*. Furthermore, the TR-FRET analysis revealed that OmKAI2d3 is activated by (+)-GR24 with affinity in the tens of nanomolar range, suggesting that the strong ligand perception ability of OmKAI2d3 is responsible for the SL hypersensitivity of *O. minor* seeds.

**Figure 4.**
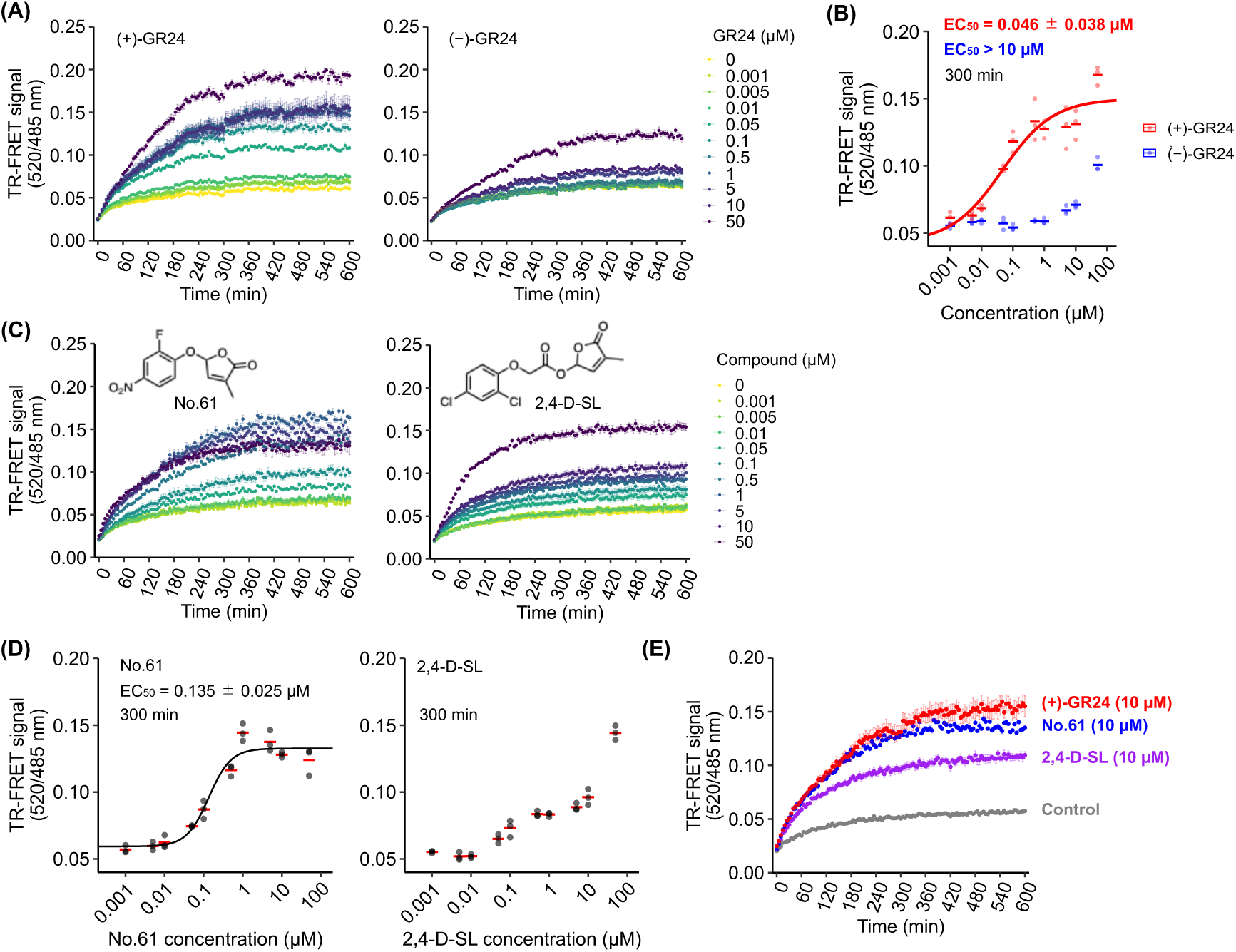
Quantitative and kinetics analyses of the *O. minor* SL signaling complex formation triggered by various agonists. (A) Kinetic measurements of OmKAI2d3–OmSMAX1-D1M TR-FRET signals in the presence of indicated concentration of (+)-GR24 and (−)-GR24. The assay was performed using OmKAI2d3-mEGFP (100 nM) and 3×FLAG-OmSMAX1-D1M (5 µg/mL). Data are the means ± SE (*n* = 3). (B) Dose-titration of (+)-GR24 and (−)-GR24 in OmKAI2d3–OmSMAX1-D1M TR-FRET assay at the single time point (300 min) extracted from the data as shown in Figure 4A. Dots represent individual data values of three technical replicates. (C) Kinetic analyses of agonistic activity of No.61 and 2,4-D-SL on OmKAI2d3–OmSMAX1-D1M complex formation. The assays were performed using OmKAI2d3-mEGFP (100 nM) and 3×FLAG-OmSMAX1-D1M (5 µg/mL). Data are the means ± SE (*n* = 3). (D) Dose-titration of No.61 and 2,4-D-SL in OmKAI2d3–OmSMAX1-D1M TR-FRET assay at the single time point (300 min) extracted from the data as shown in Figure 4C. Dots represent individual data values of three technical replicates. (E) Kinetic comparison of TR-FRET signals upon addition of the SL receptor agonist (10 µM each) extracted from Figure 4A,C.

### Kinetic analysis of the OmKAI2d3–OmSMAX1 interaction using various SL analogs

We found that the TR-FRET signal increased in time-dependent manner upon treatment with (+)-GR24 over the time scale of hours. Considering the hydrolytic activity of SL receptors, we hypothesized that the time-dependency of the SL-induced PPI signal is associated with the hydrolysis of SL agonists. To investigate the relationship between SL hydrolysis and PPI induction, we performed kinetic analyses of the PPI-inducing activities of three SL analogs, namely, (+)-GR24, No.61, and 2,4-D-SL. It has been reported that No.61 is more resistant to OmKAI2d3-mediated hydrolysis than is *rac*-GR24, and that No.61 strongly induces germination of *O. minor* (Kawada *et al*., 2024). In addition, it has been reported that 2,4-D-SL is more quickly hydrolyzed than is *rac*-GR24, and that 2,4-D-SL potently inhibits the hydrolysis of Yoshimulactone Green, a pro-fluorescent SL agonist (Tsuzuki *et al*., 2024). Despite its ability to bind to OmKAI2d3, 2,4-D-SL exhibited weaker activity to induce *O. minor* germination compared with *rac*-GR24. The TR-FRET assay showed that both No.61 and 2,4-D-SL induced the PPI of OmKAI2d3–OmSMAX1-D1M (Figure 4C). The EC50 value of No.61 was estimated to be 0.135 µM, consistent with its strong ability to induce *O. minor* germination (Figure 4D). In contrast, 2,4-D-SL did not elicit a sigmoidal dose-response curve (Figure 4D). Remarkably, we also found that maximum signal intensities differed depending on the type of SL agonist. As shown in Figure 4E, when comparing the time-dependent PPI signals among three agonists at 10 µM, the signal induced by 2,4-D-SL stopped earlier than did the signals induced by (+)-GR24 and No.61. Consequently, the maximum signal intensity induced by 2,4-D-SL was lower than those induced by (+)-GR24 and No.61 (Figure 4E). Considering its high susceptibility to OmKAI2d3-mediated hydrolysis, these results suggest that the rapid hydrolysis of 2,4-D-SL may prevent effective and stable induction of the PPI. Therefore, our TR-FRET assay system offers novel insights into PPI dynamics based on their momentary affinity, hydrolyzability, and receptor activation potential.

### Selective sensing of the naturally occurring SL receptor agonists from root exudate

As described above, the TR-FRET assay system with the hypersensitive SL receptor OmKAI2d3 easily and quantitatively detected several hundred picograms of active SL in a 20-µL reaction volume. In general, the time-gated and ratiometric measurements in TR-FRET assays suppress interference from irrelevant compounds (Bazin *et al*., 2002). Thus, we speculated that this method would be a useful tool to detect naturally occurring SLs from crude samples containing diverse natural products. Although the full-length OmSMAX1 protein was degraded as mentioned above, we detected the SL-dependent TR-FRET signal from a mixture containing the full-length OmSMAX1 and many degradation products with a slightly higher signal to noise ratio that that of a mixture containing OmSMAX1-D1M (Figure S9). Accordingly, we used 3×FLAG-OmSMAX1-6×His in SL-sensing experiments. The FRET efficiency is influenced by the donor/acceptor wavelength, orientation, and distance. Terbium-cryptate is one of the most popular lanthanide complexes for the TR-FRET assays by combining with the various types of green- and red-fluorescent dyes as the acceptors. Thus, toward further optimization of the signal to nose ratio, we adapted the SNAP tag labeled by fluorescein (FL) or sulfo-cyanine5 (Sulfo-Cy5) instead of mEGFP (Keppler *et al*., 2003). Furthermore, as the structural basis of the complex between the SL receptor and the transcriptional repressor had never been experimentally revealed, we roughly tested two different linker lengths between OmKAI2d3 and mEGFP. Even though these were not significantly optimized conditions, we demonstrated the capability of the FL- or Sulfo-Cy5-labeled SNAP tag in addition to mEGFP (Figure S10A,B). In the following SL sensing experiments, we used the pair 3×FLAG-OmSMAX1-6×His and 6×HN-OmKAI2d3-GSSG-mEGFP, in which mEGFP was connected at the C-terminus by a Gly-Ser-Ser-Gly short linker. We performed TR-FRET assays using extracts of rice (cv. Shiokari) hydroponic cultures that contained numerous organic compounds, including SLs such as 4-deoxyorobanchol (Figure 5A) (Umehara *et al*., 2008). We found that root exudates of *d14*, the rice mutant that strongly accumulates SL because it lacks the SL receptor, strongly increased the TR-FRET signal (Figure 5B). In contrast, root exudates of *d10*, the rice SL biosynthesis mutant defective in endogenous SLs, barely generated a response (Figure 5B) (Arite *et al*., 2009; Umehara *et al*., 2008; Arite *et al*., 2007). Additionally, the addition of *rac*-GR24 to *d10* root exudate restored the TR-FRET signal to the same level as that obtained with *d14* root exudate, indicating that agonistic activities can be specifically detected even in crude samples (Figure 5C). In addition, the EC50 value of (+)-GR24 was hardly influenced by the *d10* root exudate (Figure S11). These results clearly demonstrate the utility of this method for specifically detecting naturally occurring SLs with minimal interference.

**Figure 5.**
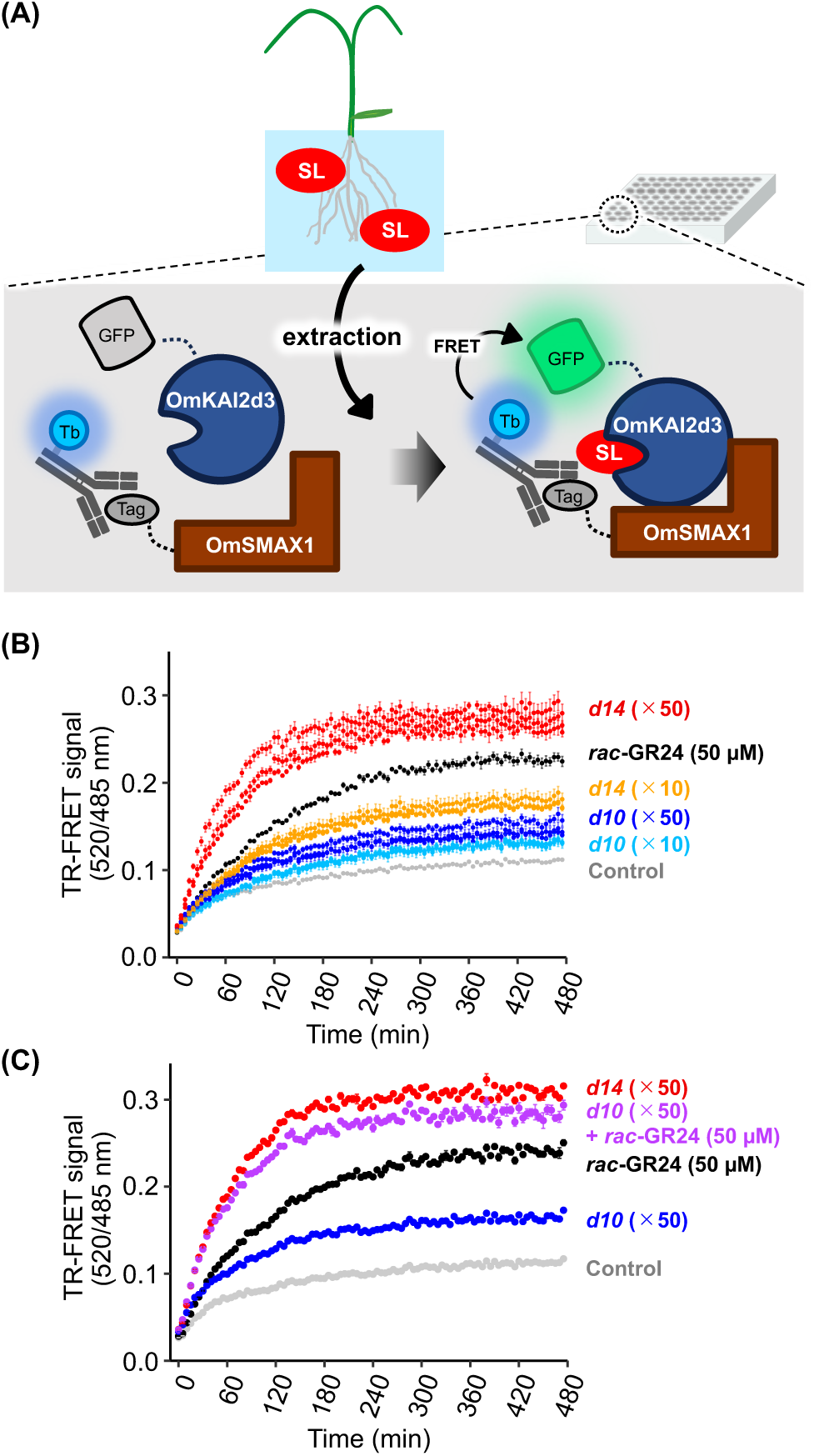
Application of the TR-FRET assay system for sensing the naturally occurring SLs. (A) Schematic of the high-throughput TR-FRET-based SL sensing system. Natural compound mixtures are extracted from root exudates and subjected to a 96-well TR-FRET assay format using the highly-sensitive OmKAI2d3–OmSMAX1 pair without further purification. (B) SL detection from root exudate of rice mutants (*d10*: SL deficient mutant, *d14*: SL accumulated mutant) using the TR-FRET system. Each data represents the individual assay values of three biological replicates. (C) Complementation of weak TR-FRET signals in *d10* root exudate by supplementing *rac*-GR24. The assays were performed using OmKAI2d3-GSSG-mEGFP (100 nM) and 3×FLAG-OmSMAX1 (5 µg/mL). Concentrations of root exudate samples are expressed as a fold enrichment relative to the original volume of the hydroponic culture media. Data are the means ± SE (*n* = 3).

## Discussion

We developed an assay system to monitor the formation of SL signaling complexes derived from *A. thaliana* and *O. minor* using TR-FRET technology. Additionally, we demonstrated that our TR-FRET assay system can distinguish the mode of action of different types of ligands for SL receptors. For example, assays with (+)-GR24 and (−)-GR24 exhibited remarkable differences in the maximum signal intensity, rather than EC50 values, of AtD14–AtSMXL7 complex formation. Considering the weak but significant activity of (−)-GR24 to inhibit branching in Arabidopsis, our results suggest that (−)-GR24 might function as a partial agonist for AtD14 (Scaffidi *et al*., 2014). It is noteworthy that our TR-FRET assay system defines the degree of agonistic activity semi-quantitatively, reflecting *in vivo* activity. Even though the mechanism of partial activation of AtD14 by (−)-GR24 is unclear, the weak induction of the TR-FRET signal suggests that the free energy landscape of AtD14 activation depends on the ligand configuration (Chen and Shukla, 2023; Chen *et al*., 2022).

Our kinetic analyses revealed that, although adding an antagonist abolished the activation of AtD14, it did not disrupt the already-formed AtD14–AtSMXL7 complex. One possibility is that the dissociation of AtD14–AtSMXL7 is quite slow because of multi-point interactions on the protein–protein contact surface. Meanwhile, mass spectrometric and crystallographic analyses of SL-supplied receptors have demonstrated that the methyl butanolide ring is covalently trapped at the catalytic amino acid residue. Therefore, it is also possible that this covalent attachment prevents the antagonist from binding. In either case, the results provide new insights into the dynamics of the SL signaling complex, as well as the importance of the timing of antagonist treatment.

The kinetic TR-FRET assay showed the signal induction by 2,4-D-SL decreased faster than that by (+)-GR24 in the OmKAI2d3–OmSMAX1-D1M complex. This observation indicates that excessively hydrolysable SL analogs cannot sufficiently activate SL receptors for complex formation, possibly because the receptor-mediated hydrolytic deactivation of these chemicals is too fast. Accordingly, although we have not yet concluded whether the PPI occurs before or during the hydrolysis process, strong tolerance of SL agonists to receptor-mediated hydrolysis would contribute to the effective activation of the SL signal.

We observed slight increases in TR-FRET signals even in the absence of agonists in the AtD14–AtSMXL7 and OmKAI2d3–OmSMAX1-D1M pairs, but not in the AtD14–MAX2 pair (Figure S2B; Figure S7). The trend was also observed in our pull-down assays (Figure 1A; Figure 3A) as well as in previous studies (Yao *et al*., 2017; Jiang *et al*., 2013). These results indicate that the receptor proteins can partially recruit the repressor proteins independently of SL. To prevent inappropriate activation of SL signaling *via* degradation of repressors, the recruitment of the E3 ubiquitin ligase that includes MAX2 is likely strictly controlled by SL. Furthermore, we should investigate whether the SL-independent receptor–repressor interaction occurs *in planta* and influences the regulation of SL signaling. This is especially important for obligate parasitic plants because tight control of the signal repression mechanism is crucial to prevent death *via* undesired germination in the absence of host plants.

The dynamics of PPIs in receptor complexes of the plant hormone auxin have been analyzed using surface plasmon resonance (SPR), providing critical insights into the molecular mechanism of auxin signaling (Calderón Villalobos *et al*., 2012). For example, different auxins differentially induce the formation of auxin co-receptor complexes, as measured by their association and dissociation rates (Calderón Villalobos *et al*., 2012). In addition, the binding kinetics differ among the auxin co-receptor pairs, which are highly duplicated in angiosperms (Calderón Villalobos *et al*., 2012). As shown in studies on auxin, information on the kinetic and thermodynamic characteristics of ligand-induced PPIs is crucial not only for understanding the biological functions of plant hormones, but also for the development of highly potent ligands. However, few studies have explored the kinetics of ligand-induced PPIs under the research of other plant hormones. TR-FRET is a promising high-throughput platform to quantitatively and kinetically characterize molecular interactions, and it is widely used in the field of drug discovery (Greenway *et al*., 2024; Falkenstern *et al*., 2024). In this study, we developed a kinetic assay system for monitoring the formation of the SL signaling complex using TR-FRET, offering a novel strategy for future research in SL signaling. In addition, our study provides a technical basis for establishing a similar kinetic assay format for other plant hormones that induce PPIs (Takeuchi *et al*., 2021).

We also demonstrated that our TR-FRET assay can selectively detect naturally occurring SLs from plant-derived samples containing numerous natural products. Compared with conventional liquid chromatography–tandem mass spectrometry and phenotype-based SL detection, the TR-FRET system has advantages of rapid and activity-based detection of SL molecules, as well as compatibility with impurities that inhibit plant growth and development. We also demonstrated the use of the SNAP tag, which is easily labeled by FL and Sulfo-Cy5, as the acceptor in our TR-FRET assay system. The multi-color platform will enable robust screening with validation of true fractions containing active compounds (Hassig *et al*., 2014). The structures of SLs are diversified among plant species, and more than 30 SL derivatives have been discovered (Chesterfield *et al*., 2020). However, the active hormone in the shoot-branching pathway has not been identified (Seto, 2023). We are currently working to detect and isolate unknown active hormones related to the inhibition of shoot branching, as well as diverse natural compounds with SL-like activities that are biosynthesized in various species.

Surprisingly, we observed a fast and strong increase in the PPI signal upon addition of *d14* root exudate, compared with that obtained using synthetic *rac*-GR24 as an authentic control. Moreover, the addition of *rac*-GR24 to *d10* root exudate restored the PPI signal to the same level as that obtained with *d14* root exudate. Rather than differences in activity between GR24 and natural SLs, these results suggest that rice root exudate contains putative enhancers, such as cofactors or allosteric modulators, for the formation of the OmKAI2d3–OmSMAX1 complex. Previous crystallographic and biochemical studies indicate that auxin and jasmonate receptors require inositol polyphosphates as essential cofactors (Sheard *et al*., 2010; Tan *et al*., 2007). Likewise, in the SL signaling complex, the structural dynamics of MAX2 are controlled by citrate, a widely distributed organic acid, and the citrate-dependent conformational switch of MAX2 is thought to be associated with SL signal transduction (Tal *et al*., 2022; Shabek *et al*., 2018). Thus, it is possible that there are modulators of the SL receptor–repressor interaction, although further research is needed to identify and functionally characterize them.

Application of the TR-FRET system for other plant hormones will be useful to identify unknown chemicals that control plant hormone signaling. Notably, despite previous attempts to identify the endogenous ligand for KAI2, the chemical identity of this plant hormone candidate has not yet been discovered (Waters and Nelson, 2022). Given the high structural similarity of KAI2 to SL receptors such as OmKAI2d3 and AtD14, it will be feasible to develop a KAI2-based TR-FRET system that can be used to discover unknown KAI2 ligands.

In summary, we developed a novel method based on TR-FRET to monitor the formation of SL signaling complexes. This kinetic and high-throughput assay system will facilitate further research on the dynamics of important molecular interactions among SL signaling components. In addition, activity-based screening using this method will allow for the discovery of potent SL signaling modulators from biological resources and drug-like chemical libraries.

## Materials and Methods

### Chemicals and general methods

Tolfenamic acid was obtained from Tokyo Chemical Industry Co., Ltd. (Tokyo, Japan). (+)-GR24 and (−)-GR24 were prepared as described in supporting information. No.61, 2,4-D-SL, and the fluorescein *O*^6^-benzylguanine conjugate (FL-BG) were synthesized as described in previous papers (Kawada *et al*., 2024; Tsuzuki *et al*., 2024; Ramil *et al*., 2017). The sulfo-Cy5 *O*^6^-benzylguanine conjugate (Sulfo-Cy5-BG) was synthesized as described in supporting information.

All reagents for chemical syntheses were purchased from Tokyo Chemical Industry Co., Ltd. (Tokyo, Japan) and FUJIFILM Wako Pure Chemical Corp. (Osaka, Japan). NMR spectra were measured by a NMR spectrometer JNM-ECZL (JEOL). High-resolution mass spectrometry (HRMS) was performed by a quadrupole/time-of-flight tandem mass spectrometer X500R (AB SCIEX).

### Vector construction

To substitute the 6×His tag of pET47b to 6×HN tag, the hybridized DNA oligos coding 6×HN tag, described in Table S1, was cloned into linearized pET47b at the site between NdeI and SacⅡ using the In-Fusion cloning Kit (Takara). Hereafter, we referred to this vector as pET47b-6×HN. To prepare vectors fusing mEGFP and SNAP tag at C-terminus, each artificial DNA fragment (Eurofins Genomics) was inserted into the site between XhoI and PacI of pET47b-6×HN based on In-Fusion cloning. For the N-terminal tagging of mEGFP and SNAP, each coding sequence was amplified by PCR and individually inserted into the site between SmaI and SacI. For preparing the protein expression vectors of AtD14 and OmKAI2d3 connecting mEGFP/SNAP based on their modified pET47b-6×HN, the coding sequences of AtD14, AtD14^S97A^ and OmKAI2d3, which have been used in our previous studies (Seto *et al*., 2019; Takei *et al*., 2023), were amplified by PCR with primers described in Table S1. Each PCR product was introduced into the site between SmaI–SacI and XhoI–PacI for the N-terminal and C-terminal tagging, respectively. To substitute the original linker (the sequence of thrombin and S-tag) in the middle of OmKAI2d3 and mEGFP with Gly-Ser-Ser-Gly (GSSG) linker, the annealed DNA fragment as shown in Table 1 was cloned into the linearized pET47b-6×HN vector carrying OmKAI2d3 and mEGFP digested at SacI and XhoI sites.

The protein expression vectors for AtSMXL7 and OmSMAX1(-D1M) were constructed by using pET26b, which encodes pelB signal sequence and C-terminal 6×His tag, according to the previous study (Khosla *et al*., 2020), with slight modifications. The hybridized 3×FLAG tag-coding oligos described in Table S1 was cloned into the site between NcoI and BamHI of pET26b by In-Fusion cloning, yielding pET26b-3×FLAG. The cording sequence of AtSMXL7 (Seto *et al*., 2019) was amplified by PCR with primers described in Table S1. The cording sequence of OmSMAX1 which is predicted from genome data of *O. minor* (Bürger *et al*., 2025) was amplified from cDNA of *O. minor* by PCR with primers described in Table S1. The region of OmSMAX1-D1M (A164 to K610) was determined according to the previous study (Khosla *et al*., 2020). Each DNA fragment was cloned into the site between BamHI and XhoI of pET26b-3×FLAG by In-Fusion cloning. For the construction of StrepⅡ-HiBiT-AtSMXL7-6×His protein expression vector, the insert DNA fragment of StrepⅡ-HiBiT was prepared by primer extension using partially complemental oligos described in Table S1. The elongated DNA fragment was cloned into pET26b by In-Fusion cloning. The following operations were carried out as described above.

The CDSs of AtMAX2 and ASK1 (Seto *et al*., 2019) were amplified by PCR with primers described in Table S1. For the transient co-expression of 6×His-AtMAX2 and tag-free ASK1 in tobacco leaves, these PCR products were sub-cloned into pDONR207 by BP reaction and recombined into pEAQ-HT DEST 2 and pEAQ-HT DEST1 for AtMAX2 and ASK1, respectively (Sainsbury *et al*., 2009). For 6×His-3×FLAG-AtMAX2 expression, the plasmid was prepared following GoldenBraid assembly systems previously developed (Sarrion-Perdigones *et al*., 2013). The GoldenBraid 2.0 kit was a gift from Diego Orzaez (Addgene kit # 1000000076). The entry vectors of 6×His-3×FLAG for insertion at B3 site and of AtMAX2 for that at B4–B5 site were sub-cloned into pUPD2 vector via Golden Gate cloning by using BsmBI restriction enzyme and DNA ligase (New England Biolabs). Then, sequences of Arabidopsis Ubiquitin10 promoter (GoldenBraid 2.0 kit), 6×His-3×FLAG, AtMAX2, and Arabidopsis Ubiquitin3 terminator (GoldenBraid 2.0 kit) within pUPD2 were assembled into pDGB3 vector by BsaI restriction enzyme and DNA ligase (New England Biolabs).

Protein expression and purification

AtD14, AtD14^S97A^ and OmKAI2d3 protein connecting mEGFP or SNAP were purified as previously described and stored at –80°C until use (Tsuzuki *et al*., 2024).

The expression and purification of AtSMXL7 and OmSMAX1 proteins was performed according to the previous study (Khosla *et al*., 2020) with slight modifications. After protein expression induction, the *E. coli* cells were harvested by centrifugation at 3,700 × g and the pellet were stored at −30℃ until use. The pellet was resuspended in lysis buffer [50 mM sodium phosphate buffer (pH 7.6), 300 mM NaCl, 5 mM imidazole, 1× EDTA-free protease inhibitor cocktail (nacali tesque), 1.5 mg/mL Lysozyme, 0.1% Tween 20, with or without 10 unit/mL Benzonase (Merck)] and incubated for 30–60 min on ice. The resuspended cells were lysed with gentle sonication. After centrifugation at 18,000 × g for 30 min, the supernatant was subjected affinity chromatography using nickel sepharose resin. The column was washed with the washing buffer [50 mM sodium phosphate buffer (pH 7.6), 300 mM NaCl, 5–20 mM imidazole]. The bound protein was eluted with the elution buffer [50 mM sodium phosphate buffer (pH 7.6), 300 mM NaCl, 250 mM imidazole]. The eluate was concentrated using VIVASPIN Turvo15 (Sartorius) and buffer-exchanged into 1× PBS (–). The protein solution was aliquoted to the appropriate volume, immediately frozen in liquid nitrogen, and stored at −80℃ until use. OmSMAX1-D1M was prepared by the same procedure as described above with slight changes in buffers. Lysis buffer [50 mM sodium phosphate buffer (pH 7.0), 500 mM NaCl, 1× EDTA-free protease inhibitor cocktail, 1.5 mg/mL Lysozyme, 0.1% Tween 20, 10 unit/mL Benzonase], washing buffer [50 mM sodium phosphate buffer (pH 7.0), 300 mM

NaCl, 20 mM imidazole], and elution buffer [50 mM sodium phosphate buffer (pH 7.0), 300 mM NaCl, 200 mM imidazole] were employed for OmSMAX1-D1M purification.

For the expression of AtMAX2 protein, the expression vectors containing AtMAX2 gene and ASK1 were introduced into *Agrobacterium tumefacience* LBA4404, respectively. The transformed Agrobacterium cells were pre-cultured in LB selection medium at 30℃ with shaking. Overnight cultures were diluted into fresh LB medium and incubated at 30℃ until the OD600 reached to 0.5–1.0. The Agrobacterium cells were harvested by centrifugation at 3,500 × g and re-suspended in infiltration buffer [10 mM MES-KOH (pH 5.6), 150 mM acetosyringone, 10 mM MgCl2] to adjust OD600 to 0.5-1.0. After placing under dark conditions for at least 1 h, the Agrobacterium solutions for AtMAX2 and ASK1 expression were mixed (1:1) and infiltrated into *N. benthamiana* leaves. The leaves were harvested at 5-7 days post-infiltration. 10 g of the leaves were ground in a mortar with the aid of liquid nitrogen. The powdered leaves were extracted with 15 mL of the extraction buffer [50 mM HEPES-NaOH (pH 8.0), 150 mM NaCl, 5 mM 2-mercaptoethanol, 50 µM MG-132, 10% glycerol, 0.5% Triton X-100, 1× EDTA-free protease inhibitor cocktail]. After the centrifugation at 18,000 × g for 15 min, the supernatants were filtered through Miracloth (Merck). The filtrate was centrifuged at 18,000 × g for 15 min. The supernatant was subjected affinity chromatography using TALON superflow resin (Cytiva). The column was washed with the washing buffer [50 mM HEPES-NaOH (pH 8.0), 150 mM NaCl, 5 mM imidazole, 0.5% Triton X-100). The bound protein was eluted with the 1×PBS (–) containing 150 mM imidazole. The eluate was concentrated using VIVASPIN Turvo15 (Sartorius) and buffer-exchanged into 1× PBS (–). The protein solution was aliquoted to the appropriate volume, immediately frozen in liquid nitrogen, and stored at −80℃ until use.

### Fluorescent labelling of SNAP tag

SNAP-tagged proteins (50 µM) were incubated with FL-BG (200 µM) or Sulfo-Cy5-BG (200 µM) in 1× PBS (–) containing 1mM DTT at 25℃ for 2 h. Un-reacted labelling regents were removed by using desalting column (PD-10; Cytiva). After concentrating protein solutions by VIVASPIN, the concentration of each protein was determined by the Bradford method. The labeling yield were calculated from the absorbance (FL: the molar extinction coefficient of 68,000 M^−1^cm^−1^ at 494 nm; Sulfo-Cy5: the molar extinction coefficient of 250,000 M^−1^cm^−1^ at 649 nm) and estimated as >70%.

### *In vitro* Pull-down assay

AtD14-mEGFP (25 µg) was immobilized on 10 µL of GFP-Trap Magnetic Particles M-270 (Proteintech) at 4℃ for 1 h with agitation. The beads were washed with 500 µL of washing buffer [50 mM MES-NaOH (pH6.5), 150 mM NaCl, 0.1 mg/mL BSA, 10% glycerol, 0.1 % Tween 20] for 3 times. AtSMXL7 protein (45 µg) or AtMAX2 protein (20 µg) was added to the beads in 40 µL of assay buffer [50 mM MES-NaOH (pH6.5), 150 mM NaCl, 2 mM DTT, 0.1 mg/mL BSA, 10% glycerol, 1% DMSO, 0.1 % Tween 20] with or without 50 µM GR24. After incubation at room temperature for 1 h with agitation, the beads were washed by 200 µL of washing buffer for 5 times. Bound proteins were extracted by 20 µL of 1×SDS-sample buffer with boiling. The samples were subjected to SDS-PAGE and western blotting analysis.

OmKAI2d3-mEGFP (12.6 µg) was immobilized on 10 µL of GFP-Trap Magnetic Particles M-270 at 4℃ for 1 h with agitation. The beads were washed with 500 µL of washing buffer [50 mM MES-NaOH (pH6.5), 150 mM NaCl, 0.1 mg/mL BSA, 10% glycerol, 0.1 % Tween 20] for 3 times. OmSMAX1-D1M protein (0.75 µg) was added to the beads in 40 µL of assay buffer [50 mM MES-NaOH (pH6.5), 150 mM NaCl, 2 mM DTT, 0.1 mg/mL BSA, 10% glycerol, 1% DMSO, 0.1% Tween 20] with or without 50 µM *rac*-GR24. After incubation at 4℃ for 30 min with agitation, the beads were washed by 200 µL of washing buffer for 5 times. Bound proteins were extracted by 20 µL of 1×SDS-PAGE sample buffer with boiling. The samples were subjected to SDS-PAGE and western blotting analysis.

### Western blotting

Protein samples were separated by SDS-PAGE and transferred to a nitrocellulose membrane by semi-dry blotting system (ATTO). For the FLAG-tagged, His-tagged and mEGFP-tagged proteins, the membrane was blocked in 1×TBS-T containing 0.5% BSA at room temperature for 1 h. These proteins were detected by using 1:2000 dilution anti-DDDDK (FLAG)-tag monoclonal antibody-HRP conjugate (Medical & Biological Laboratories), 1:5000 anti-6×His monoclonal antibody-HRP conjugate (FUJIFILM Wako Pure Chemical) and 1:2000 dilution anti-GFP polyclonal antibody-Alexa Fluor 488 conjugate (Invitrogen). Chemiluminescent detections were carried out using EzWestLumi one (ATTO) and images were obtained using the iBright CL 1500 (Invitrogen). The HiBiT-tagged AtSMXL7 was detected by using Nano-Glo HiBiT Blotting System (Promega) according to manufactures protocol.

### TR-FRET assay

The experiments were performed in HTRF 96-well low-volume white plate (Revvity) in 20 µL of reaction buffer [50 mM MES-NaOH (pH6.5), 150 mM NaCl, 2 mM DTT, 0.1 mg/mL BSA, 1× HTRF mAb anti-FLAG Tb-Conjugate (Revvity)] with the indicated concentrations of proteins, chemicals and solvents (acetone or DMSO). Unless otherwise stated, the final concentration of acetone was adjusted to 1% (v/v). After the FLAG-tagged protein was incubated in reaction buffer at room temperature for 30 min, the SL receptor proteins and chemical solutions were added to a reaction mixture on ice. Measurements were taken on a microplate reader Infinite 200 PRO F Plex (TECAN) with the following settings (mEGFP and SNAP-FL: excitation 340(35)nm, emission A 520(10)nm, emission B 485(20)nm, 120 µs delay time, 200 µs integration time, 5 min interval; SNAP-Sulfo-Cy5: excitation 340(35)nm, emission A 665(8)nm, emission B 620(10)nm, 100 µs delay time, 200 µs integration time, 5 min interval). The TR-FRET signal was taken as the emission A/emission B intensity ratio.

In the kinetic monitoring of the antagonist activity on the post-formed complex, 1 µL of tolfenamic acid solutions containing 5 % acetone was added to 20 µL of reaction solutions at 120 min post-PPI induction by (+)-GR24 (0.5 µM). Measurements were taken as described above.

### TR-FRET assay using rice root exudates

The strigolactone receptor mutant (*d14-1*) and biosynthesis mutant (*d10-1*) of *Oryza sativa* (rice) were used for the SL sensing experiments (Ishikawa *et al*., 2005). After growing for 24 days in a hydroponic culture system under phosphate deficient conditions as previously described (Umehara *et al*., 2015), root exudates of rice plants were extracted with ethyl acetate. The organic phase was dried over Na2SO4 and concentrated under a stream of nitrogen. The extracts were dissolved in DMSO and subjected to a TR-FRET assay. The experiments were performed in the reaction buffer containing 5% DMSO using the above procedure. When supplying the *rac*-GR24 or (+)-GR24 to *d10* root exudate, 0.5% or 1% acetone was added to all samples, respectively.

### Germination assay

*O. minor* germination assays were performed as previously described (Tsuzuki *et al*., 2024).

### DSF assay

AtD14 and OmKAI2d3 were cloned into pET47b-(+) vector and these proteins were prepared as previously described (Tsuzuki *et al*., 2024). AtD14 protein (5.5 µg) or OmKAI2d3 protein (10 µg) was dissolved in 20 µL of 1× PBS (–) buffer containing SYPRO Tangerine or SYPRO Orange (SigmaAldrich) and each SL analog with 1% (v/v) acetone on a 96-well PCR white plate (Roche). SYPRO Tangerine (×5000) was diluted to ×1.25 final concentration in a reaction mixture for AtD14. SYPRO Orange (×5,000) was diluted to ×0.625 final concentration for OmKAI2d3. The mixtures were heated from 20℃ to 95℃ with scanning the fluorescence intensity (Ex/Em: 483/640 nm for AtD14; Ex/Em: 483/610 nm for OmKAI2d3) using LightCycler480 (Roche). The first derivative curve was calculated from a plot of the fluorescence intensity against the temperature using the LightCycler480 Software.

### Data analysis and graphics

For the calculations of EC50 and IC50 values, the plots were fitted by a four-parameter logistic model using the R drc package (Ritz *et al*., 2015). Graphs were drawn using the R ggplot2 package (Wickham, 2016).

## Author contribution

T.S. performed majority of experiments in this paper with guidance from K.N. and Y.S.; Y.K., C.S. and T.I. performed part of protein preparation and TR-FRET analyses; M.B. supported cloning of *O*. *minor* gene; J.R. and M.K. prepared part of plasmids with guidance from S.H; K.F and T.A. prepared No.61; K.N. synthesized FL-BG and Sulfo-Cy5-BG; K.N. and Y.S. designed research. All authors wrote the manuscript.

## Acknowledgments

We thank Nagasawa Water Purification Plant for the corporation regarding *O*. *minor* sampling. We thank Dr. Jennifer Smith from Edanz (https://jp.edanz.com/ac) for editing a draft of this manuscript.

## Short legends for Supporting Information

Figure S1. Preparation of FLAG-tagged AtMAX2 and AtSMXL7 recombinant proteins.

Figure S2. TR-FRET signals in AtD14‒AtMAX2 and AtD14‒AtSMXL7 pairs in the absence or presence of *rac*-GR24 before normalization.

Figure S3. Negative control of TR-FRET assays using the catalytic mutant protein of AtD14 and FLAG-tag-free partner proteins.

Figure S4. Melting peaks of 6×His-AtD14 in the absence and presence of (+)-GR24 or (−)-GR24.

Figure S5. Phylogenetic tree of SMAX family.

Figure S6. Preparation of FLAG-tagged OmSMAX1 and OmSMAX1-D1M recombinant proteins.

Figure S7. TR-FRET signals in OmKAI2d3‒OmSMAX1-D1M pairs in the absence and presence of *rac*-GR24 before normalization.

Figure S8. Effects of (+)-GR24 and (−)-GR24 on *O. minor* seed germination and the melting temperature of OmKAI2d3.

Figure S9. TR-FRET assay using OmKAI2d3 and full-length OmSMAX1.

Figure S10. Effects of fluorophore and linker variations on the SL-dependent TR-FRET signals.

Figure S11. Effects of *d10* root exudates on the EC50 value of (+)-GR24. Preparation of (+)-GR24 and (−)-GR24. Chemical synthesis of Sulfo-Cy5-BG. Table S1. Primers used in this study.

