## Supplemental information for "*In Vitro* Dynamic and Quantitative Monitoring of Strigolactone-signaling Complex Formation by Time-resolved FRET"

### Title

**Figure S1. Preparation of FLAG-tagged AtMAX2 and AtSMXL7 recombinant proteins.**

**Figure S2. TR-FRET signals in AtD14–AtMAX2 and AtD14–AtSMXL7 pairs in the absence or presence of *rac*-GR24 before normalization.**

**Figure S3. Negative control of TR-FRET assays using the catalytic mutant protein of AtD14 and FLAG-tag-free partner proteins.**

**Figure S4. Melting peaks of 6×His-AtD14 in the absence and presence of (+)-GR24 or (−)-GR24.**

**Figure S5. Phylogenetic tree of SMAX family.**

**Figure S6. Preparation of FLAG-tagged OmSMAX1 and OmSMAX1-D1M recombinant proteins.**

**Figure S7. TR-FRET signals in OmKAI2d3–OmSMAX1-D1M pairs in the absence and presence of *rac*-GR24 before normalization.**

**Figure S8. Effects of (+)-GR24 and (−)-GR24 on *O. minor* seed germination and the melting temperature of OmKAI2d3.**

**Figure S9. TR-FRET assay using OmKAI2d3 and full-length OmSMAX1.**

**Figure S10. Effects of fluorophore and linker variations on the SL-dependent TR-FRET signals.**

**Figure S11. Effects of *d10* root exudates on the $EC_{50}$ value of (+)-GR24.**

**Preparation of (+)-GR24 and (−)-GR24.**

**Chemical synthesis of Sulfo-Cy5-BG.**

**Table S1. Primers used in this study.**

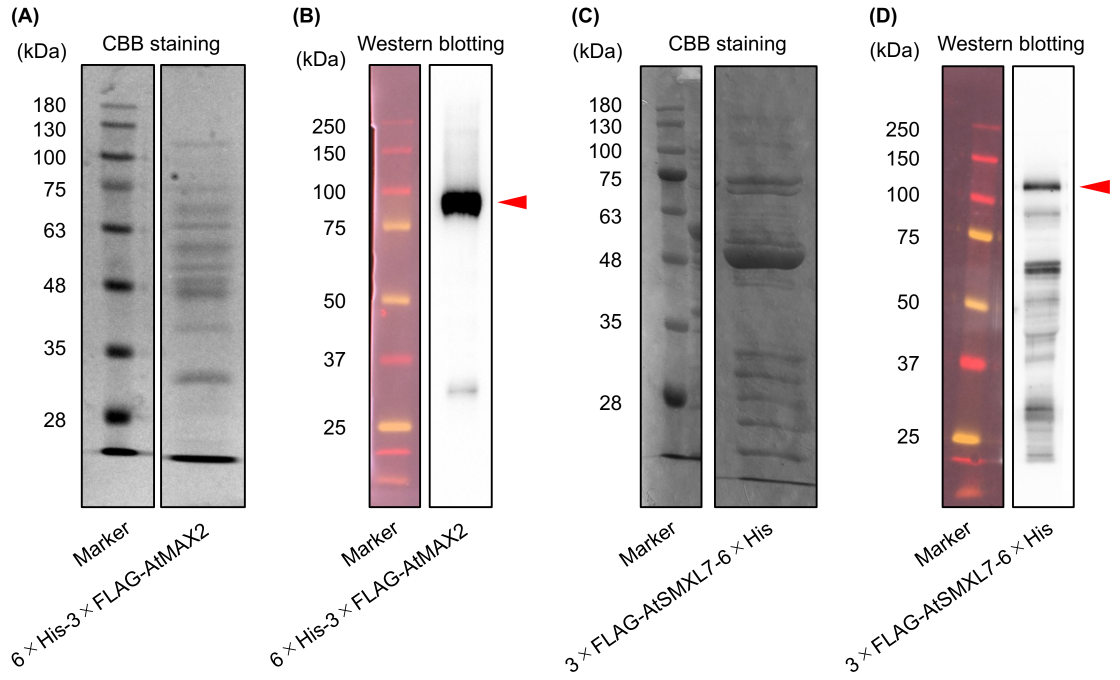


Figure S1. Preparation of FLAG-tagged AtMAX2 and AtSMXL7 recombinant proteins. (A) SDS-PAGE analysis with Coomassie Brilliant Blue (CBB) staining and (B) Western blotting analysis with anti-FLAG antibody of 6×His-3×FLAG-AtMAX2 protein after purification using TALON resin. (C) SDS-PAGE analysis with CBB staining and (D) Western blotting analysis with anti-FLAG antibody of 3×FLAG-AtSMXL7-6×His protein after purification using nickel resin. The red arrows indicate the gel bands corresponding to the size of each recombinant protein.

**(A)**

Control

| AtMAX2 (µg/mL) | - | AtD14-mEGFP (nM) 100 | AtD14-mEGFP (nM) 500 | mEGFP-AtD14 (nM) 100 | mEGFP-AtD14 (nM) 500 |
|---|---|---|---|---|---|
| - | 0.013 ± 0.0004 | 0.017 ± 0.0006 | 0.029 ± 0.0006 | 0.017 ± 0.001 | 0.027 ± 0.001 |
| 1 | 0.013 ± 0.0009 | 0.015 ± 0.0004 | 0.029 ± 0.0005 | 0.016 ± 0.0002 | 0.027 ± 0.0002 |
| 5 | 0.014 ± 0.0007 | 0.016 ± 0.0004 | 0.028 ± 0.0006 | 0.017 ± 0.0002 | 0.028 ± 0.001 |
| 10 | 0.014 ± 0.0003 | 0.017 ± 0.0006 | 0.027 ± 0.001 | 0.017 ± 0.0008 | 0.028 ± 0.0004 |

*rac*-GR24 (50 µM)

| AtMAX2 (µg/mL) | - | AtD14-mEGFP (nM) 100 | AtD14-mEGFP (nM) 500 | mEGFP-AtD14 (nM) 100 | mEGFP-AtD14 (nM) 500 |
|---|---|---|---|---|---|
| - | 0.014 ± 0.0001 | 0.020 ± 0.0009 | 0.051 ± 0.0049 | 0.022 ± 0.0011 | 0.036 ± 0.0013 |
| 1 | 0.016 ± 0.0023 | 0.021 ± 0.0009 | 0.045 ± 0.0053 | 0.021 ± 0.001 | 0.035 ± 0.0012 |
| 5 | 0.015 ± 0.0006 | 0.022 ± 0.0004 | 0.047 ± 0.004 | 0.023 ± 0.0028 | 0.038 ± 0.0026 |
| 10 | 0.016 ± 0.0013 | 0.026 ± 0.0005 | 0.059 ± 0.0078 | 0.022 ± 0.0015 | 0.038 ± 0.0063 |

TR-FRET signal (520/485 nm)
0.01 – 0.06

**(B)**

Control

| AtSMXL7 (µg/mL) | - | AtD14-mEGFP (nM) 100 | AtD14-mEGFP (nM) 500 | mEGFP-AtD14 (nM) 100 | mEGFP-AtD14 (nM) 500 |
|---|---|---|---|---|---|
| - | 0.012 ± 0.0006 | 0.015 ± 0.0002 | 0.028 ± 0.0005 | 0.015 ± 0.0003 | 0.025 ± 0.0014 |
| 1 | 0.012 ± 0.0003 | 0.020 ± 0.0008 | 0.046 ± 0.0018 | 0.020 ± 0.0004 | 0.034 ± 0.0008 |
| 5 | 0.012 ± 0.0002 | 0.025 ± 0.0022 | 0.064 ± 0.002 | 0.026 ± 0.0018 | 0.055 ± 0.0025 |
| 10 | 0.015 ± 0.0011 | 0.029 ± 0.0017 | 0.076 ± 0.0029 | 0.027 ± 0.0013 | 0.057 ± 0.0015 |

*rac*-GR24 (50 µM)

| AtSMXL7 (µg/mL) | - | AtD14-mEGFP (nM) 100 | AtD14-mEGFP (nM) 500 | mEGFP-AtD14 (nM) 100 | mEGFP-AtD14 (nM) 500 |
|---|---|---|---|---|---|
| - | 0.012 ± 0.0005 | 0.017 ± 0.0004 | 0.043 ± 0.0024 | 0.016 ± 0.0011 | 0.026 ± 0.0003 |
| 1 | 0.014 ± 0.0005 | 0.034 ± 0.0023 | 0.128 ± 0.0089 | 0.023 ± 0.0002 | 0.040 ± 0.0013 |
| 5 | 0.013 ± 0.001 | 0.062 ± 0.0043 | 0.212 ± 0.0163 | 0.037 ± 0.0011 | 0.086 ± 0.0034 |
| 10 | 0.014 ± 0.0001 | 0.076 ± 0.0098 | 0.271 ± 0.0344 | 0.036 ± 0.0007 | 0.090 ± 0.0052 |

TR-FRET signal (520/485 nm)
0.01 – 0.3

Figure S2. Values of TR-FRET signals with AtD14–AtMAX2 pair (A) and AtD14–AtSMXL7 pair (B) in the absence or presence of *rac*-GR24 at a 300 min incubation period. Data are the means ± SD ($n$ = 3).

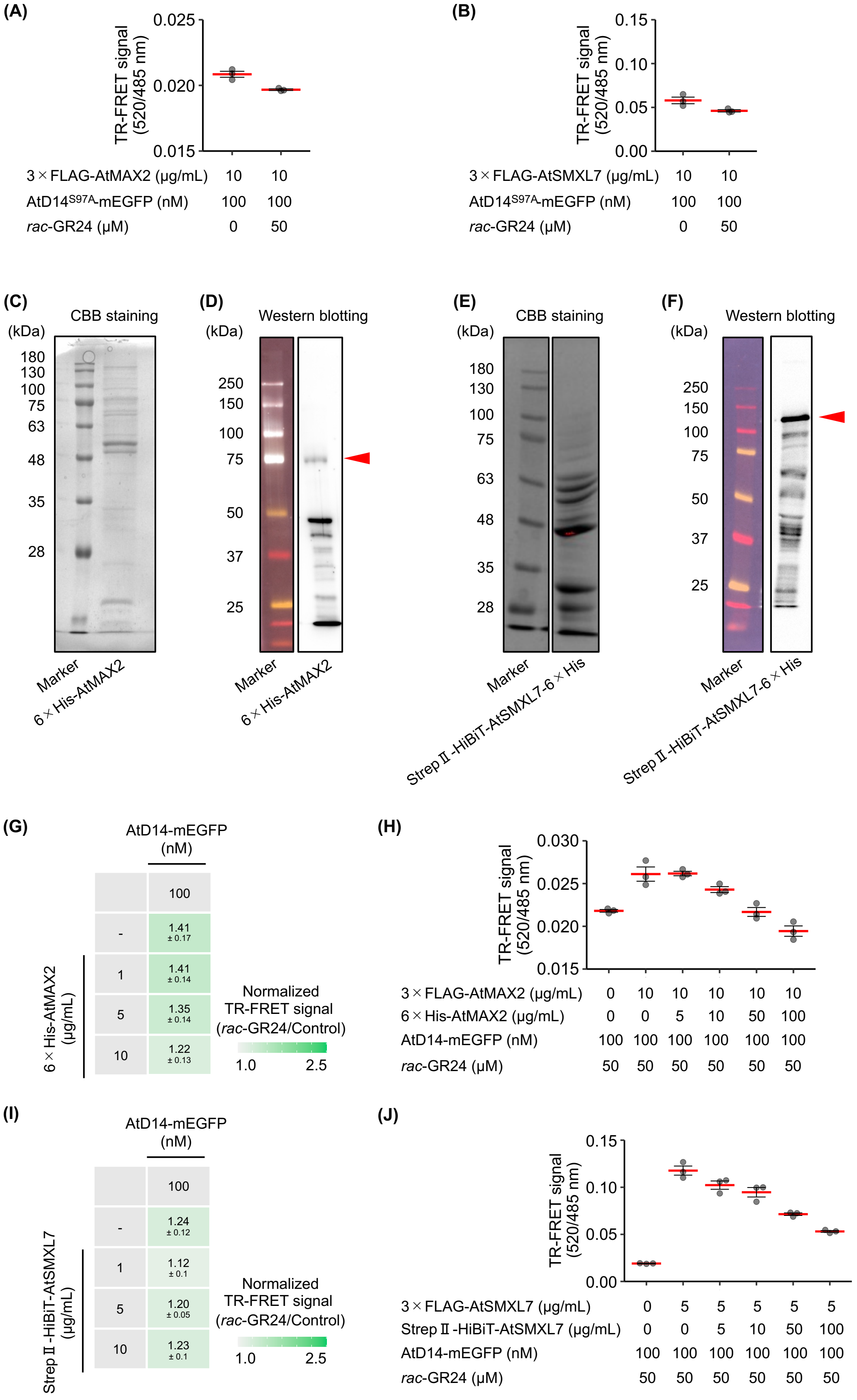

(A)
TR-FRET signal (520/485 nm)
3×FLAG-AtMAX2 (μg/mL) 10 10
AtD14S97A-mEGFP (nM) 100 100
rac-GR24 (μM) 0 50
(B)
TR-FRET signal (520/485 nm)
3×FLAG-AtSMXL7 (μg/mL) 10 10
AtD14S97A-mEGFP (nM) 100 100
rac-GR24 (μM) 0 50
(C) CBB staining
(kDa)
Marker
6×His-AtMAX2
(D) Western blotting
(kDa)
Marker
6×His-AtMAX2
(E) CBB staining
(kDa)
Marker
StrepⅡ-HiBiT-AtSMXL7-6×His
(F) Western blotting
(kDa)
Marker
StrepⅡ-HiBiT-AtSMXL7-6×His
(G)
AtD14-mEGFP (nM)
100
- 1.41 ± 0.17
1 1.41 ± 0.14
5 1.35 ± 0.14
10 1.22 ± 0.13
6×His-AtMAX2 (μg/mL)
Normalized TR-FRET signal (rac-GR24/Control)
1.0 2.5
(H)
TR-FRET signal (520/485 nm)
3×FLAG-AtMAX2 (μg/mL) 0 10 10 10 10 10
6×His-AtMAX2 (μg/mL) 0 0 5 10 50 100
AtD14-mEGFP (nM) 100 100 100 100 100 100
rac-GR24 (μM) 50 50 50 50 50 50
(I)
AtD14-mEGFP (nM)
100
- 1.24 ± 0.12
1 1.12 ± 0.1
5 1.20 ± 0.05
10 1.23 ± 0.1
StrepⅡ-HiBiT-AtSMXL7 (μg/mL)
Normalized TR-FRET signal (rac-GR24/Control)
1.0 2.5
(J)
TR-FRET signal (520/485 nm)
3×FLAG-AtSMXL7 (μg/mL) 0 5 5 5 5 5
StrepⅡ-HiBiT-AtSMXL7 (μg/mL) 0 0 5 10 50 100
AtD14-mEGFP (nM) 100 100 100 100 100 100
rac-GR24 (μM) 50 50 50 50 50 50

Figure S3. Negative control of TR-FRET assays using the catalytic mutant protein of AtD14 and FLAG-tag-free partner proteins. (A, B) TR-FRET assay using AtD14$^{S97A}$ catalytic mutant protein with 6×His-3×FLAG-AtMAX2 (A) or 3×FLAG-AtSMXL7-6×His (B). (C) SDS-PAGE analysis with CBB staining and (D) Western blotting analysis with anti-6×His antibody of 6×His-AtMAX2 protein after purification using TALON resin. (E) SDS-PAGE analysis with CBB staining and (F) HiBiT blotting using Nano-Gro HiBiT Blotting system (Promega) of StrepⅡ-HiBiT-AtSMXL7-6×His protein after purification using nickel resin. The red arrows indicate the gel bands corresponding to the size of each recombinant protein. (G) TR-FRET assay using 6×His-AtMAX2 in the presence of Tb-cryptate-modified anti-FLAG antibody. Heatmap-projected values represent fold changes of TR-FRET signal (means ± SD) upon treatment with *rac*-GR24 (50 μM) at a 300 min incubation period. (H) Anti-FLAG-based TR-FRET competition assay between 6×His-3×FLAG-AtMAX2 and 6×His-AtMAX2 in the presence of *rac*-GR24 (50 μM). (I) TR-FRET assay using StrepⅡ-HiBiT-AtSMXL7-6×His in the presence of Tb-cryptate-modified anti-FLAG antibody. Heatmap-projected values represent fold changes of TR-FRET signal (means ± SD) upon treatment with *rac*-GR24 (50 μM) at a 300 min incubation period. (J) Anti-FLAG-based TR-FRET competition assay between 3×FLAG-AtSMXL7-6×His and StrepⅡ-HiBiT-AtSMXL7-6×His in the presence of *rac*-GR24 (50 μM). (A, B, H, J) Data are the means ± SE ($n$ = 3) at a 300 min incubation period. Dots represent individual data values of three technical replicate.

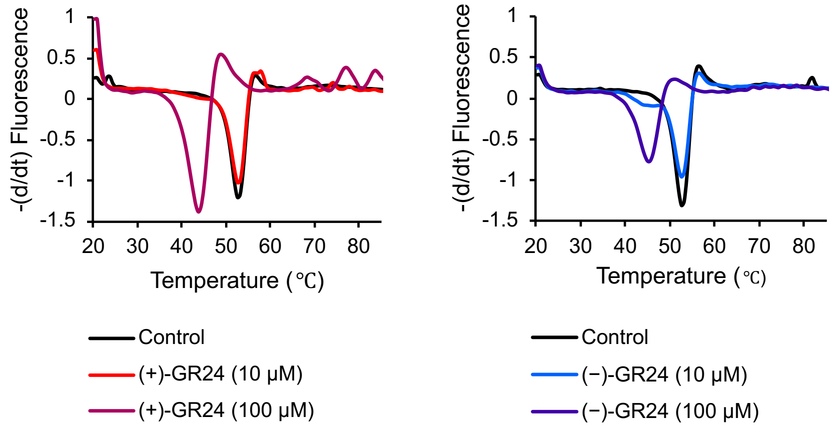


Figure S4. Melting peaks of 6×His-AtD14 in the absence and presence of (+)-GR24 or (−)-GR24. The reproducibility of the experiments was tested by making triplicate measurements.

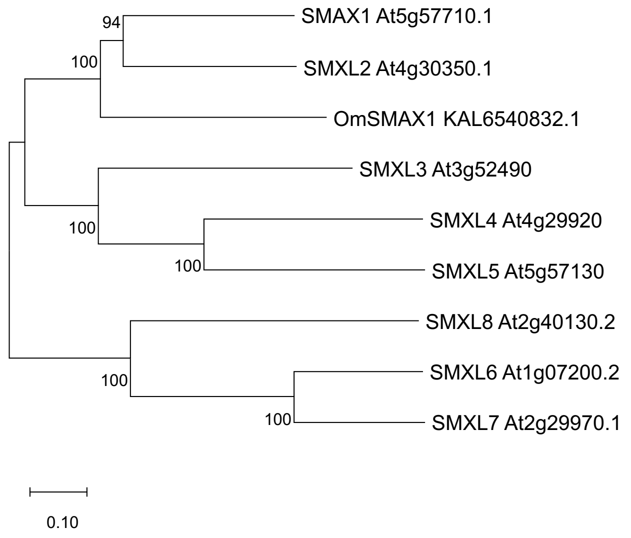


Figure S5. Phylogenetic analysis of the SMAX1-like gene from *O. minor* and SMAX1/SMXL family from *A. thaliana*. The phylogenetic tree was generated by the neighbor-joining method (bootstrap value, 1000) in MEGA11 package from the alignment of amino acid sequences of each gene product.

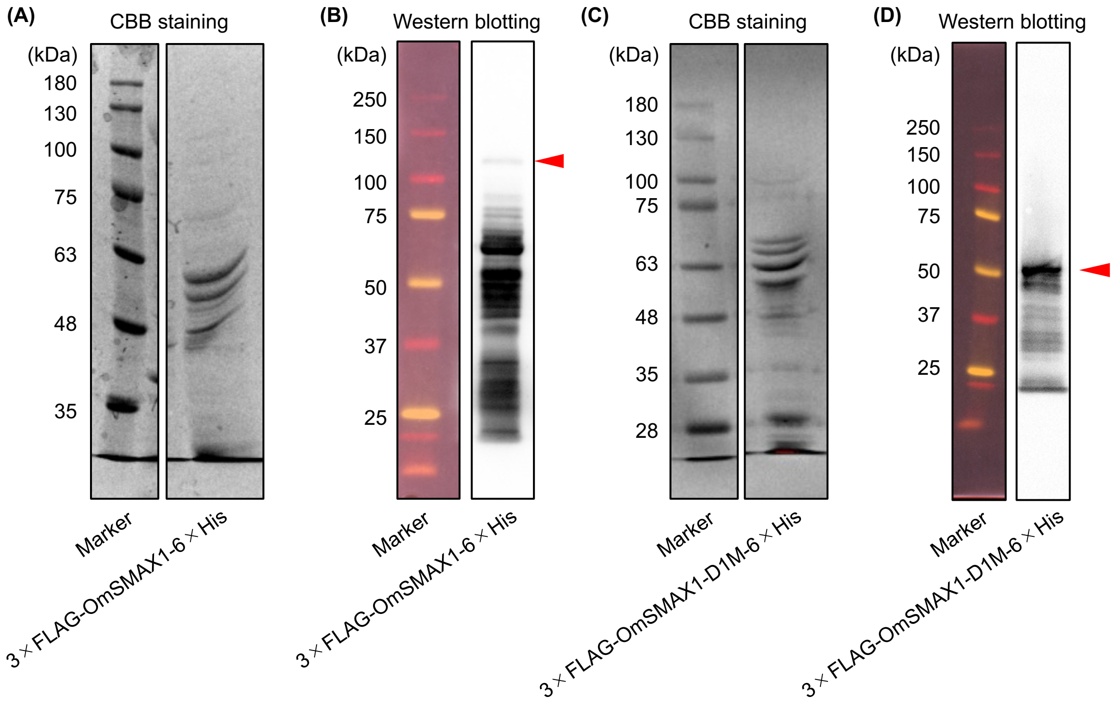


Figure S6. Preparation of FLAG-tagged OmSMAX1 and OmSMAX1-D1M. (A,C) CBB-staining SDS-PAGE and (B,D) Anti-FLAG-based western blotting analyses of 3×FLAG-OmSMAX1-6×His full-length (A,B) and D1M domain (C,D), respectively, after purification using nickel resin. The red arrows indicate the gel bands corresponding to the size of each recombinant protein.

**Control**

| OmSMAX1-D1M (μg/mL) | OmKAI2d3-mEGFP (nM): - | 100 | 500 |
|---|---|---|---|
| - | 0.011 ± 0.0004 | 0.016 ± 0.0002 | 0.033 ± 0.0003 |
| 1 | 0.013 ± 0.0003 | 0.049 ± 0.0037 | 0.089 ± 0.0011 |
| 5 | 0.013 ± 0.0005 | 0.037 ± 0.0024 | 0.097 ± 0.0011 |
| 10 | 0.014 ± 0.0001 | 0.038 ± 0.0021 | 0.103 ± 0.005 |

***rac*-GR24 (50 μM)**

| OmSMAX1-D1M (μg/mL) | OmKAI2d3-mEGFP (nM): - | 100 | 500 |
|---|---|---|---|
| - | 0.012 ± 0.0004 | 0.021 ± 0.0011 | 0.066 ± 0.0014 |
| 1 | 0.013 ± 0.0003 | 0.107 ± 0.0049 | 0.193 ± 0.0012 |
| 5 | 0.014 ± 0.0006 | 0.088 ± 0.0046 | 0.264 ± 0.0037 |
| 10 | 0.014 ± 0.0001 | 0.079 ± 0.0043 | 0.255 ± 0.0015 |

TR-FRET signal (520/485 nm): 0.01 – 0.3

Figure S7. Values of OmKAI2d3–OmSMAX1-D1M TR-FRET signals in the absence or presence of *rac*-GR24. Heatmap-projected values represent TR-FRET signal at a 300 min incubation period. Data are the means ± SD ($n$ = 3).

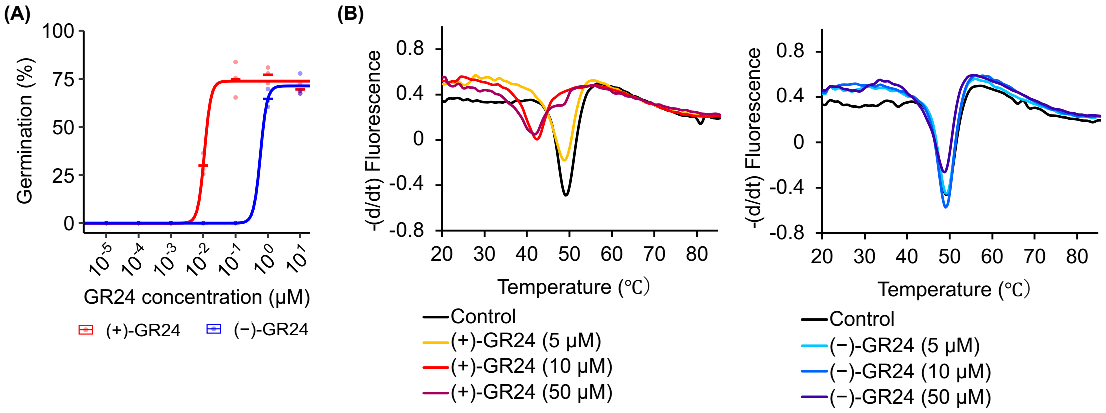


Figure S8. Effects of (+)-GR24 and (−)-GR24 on *O. minor* seed germination and the melting temperature of OmKAI2d3. (A) Germination rate of *O. minor* seeds upon addition of (+)-GR24 or (−)-GR24, measured after 5 days post-germination induction. Dots represent individual data values of three biological replicate. (B) The melting peaks of OmKAI2d3 in the absence or presence of (+)-GR24 or (−)-GR24. The reproducibility of the experiments was tested by making triplicate measurements..

| OmSMAX1 (μg/mL) | OmKAI2d3-mEGFP (nM) | |
|---|---|---|
| | - | 100 |
| - | 1.07 ± 0.08 | 1.31 ± 0.08 |
| 1 | 1.03 ± 0.03 | 2.58 ± 0.25 |
| 5 | 1.06 ± 0.02 | 2.75 ± 0.11 |
| 10 | 1.12 ± 0.05 | 2.73 ± 0.25 |

Normalized TR-FRET signal (*rac*-GR24/Control)

Low High

Figure S9. TR-FRET assay using OmKAI2d3-mEGFP and full-length OmSMAX1. Heatmap-projected values represent fold changes of TR-FRET signal upon treatment with *rac*-GR24 (50 μM) at a 300 min incubation period. Data are the means ± SD ($n$ = 3). The data in the absence of OmSMAX1 are identical to those in Figure 3B and are shown here for comparison.

**(A)**

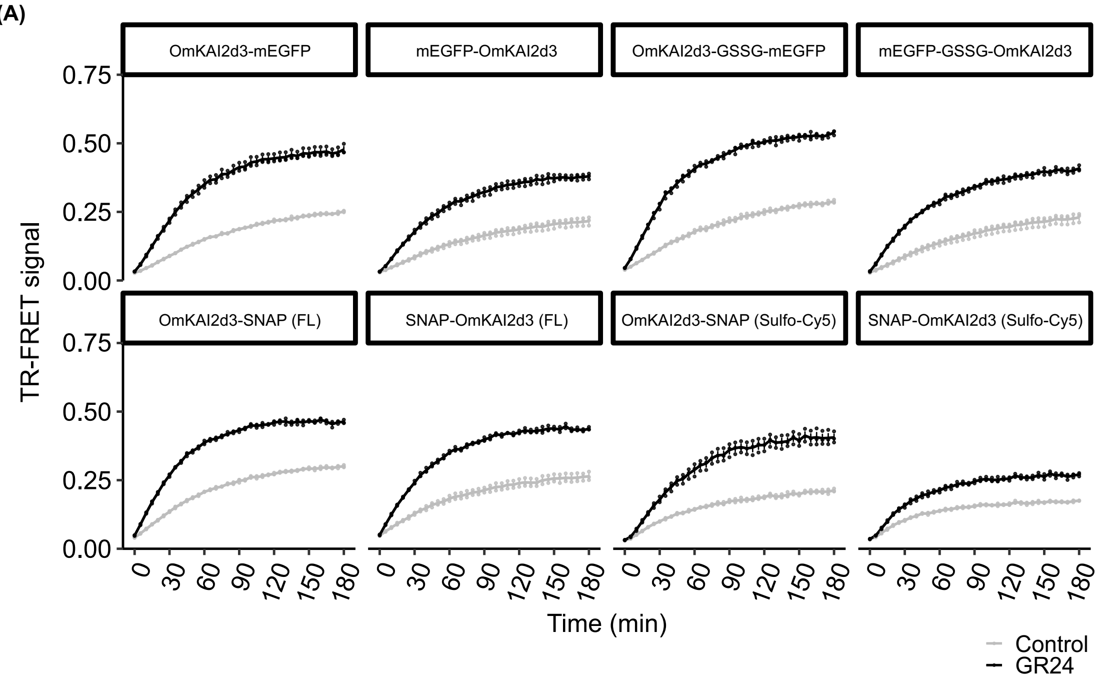


**(B)**

| | 3xFLAG-OmSMAX1 (5 μg/mL) |
|---|---|
| OmKAI2d3-mEGFP (100 nM) | 2.33 ± 0.12 |
| mEGFP-OmKAI2d3 (100 nM) | 2.06 ± 0.2 |
| OmKAI2d3-GSSG-mEGFP (100 nM) | 2.28 ± 0.13 |
| mEGFP-GSSG-OmKAI2d3 (100 nM) | 2.09 ± 0.2 |
| OmKAI2d3-SNAP (FL) (100 nM) | 1.87 ± 0.04 |
| SNAP-OmKAI2d3 (FL) (100 nM) | 1.92 ± 0.13 |
| OmKAI2d3-SNAP (Sulfo-Cy5) (100 nM) | 2.01 ± 0.19 |
| SNAP-OmKAI2d3 (Sulfo-Cy5) (100 nM) | 1.54 ± 0.07 |

Normalized TR-FRET signal (GR24/Control)

Low High

Figure S10. Effects of fluorophore and linker variations on the SL-dependent TR-FRET signals. (A) Time-course TR-FRET assay using OmKAI2d3 proteins containing indicated linkers and fluorophores. The assays were performed using fluorescently labeled OmKAI2d3 (100 nM) and 3×FLAG-OmSMAX1 (5 μg/mL). Data are the means ± SE ($n$ = 3). Dots represent individual data values of three technical replicates. (B) Heatmap-projected values represent fold changes of TR-FRET signal upon treatment with *rac*-GR24 (50 μM). Data at

60 min were extracted from the time-course shown in Figure S10A. Data are the means ± SD (*n* = 3).

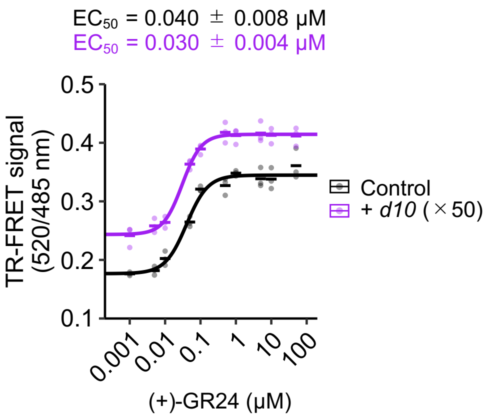


Figure S11. Effects of *d10* root exudates on the $EC_{50}$ value of (+)-GR24. Dose-titration of (+)-GR24 in OmKAI2d3-OmSMAX1 TR-FRET assay at the single time point (300 min). The assays were performed using OmKAI2d3-GSSG-mEGFP (100 nM) and 3×FLAG-OmSMAX1 (5 μg/mL). Concentrations of root exudate samples are expressed as a fold enrichment relative to the original volume of the hydroponic culture media. Control was performed without root exudates. Dots represent individual data values of three technical replicates.

**Preparation of (+)-GR24 and (−)-GR24.**

(+)-GR24 and (−)-GR24 were separated from commercially available *rac*-GR24 using HPLC equipped with CHIRALPAK IC (DAICEL, ϕ2.7×100 mm, 1.6 μm). The separation was carried out under isocratic elution using 100% MeOH at a flow rate of 2 mL $min^{-1}$. Optical rotations were measured with a JASCO DIP-370 polarimeter.

Chemical synthesis of Sulfo-Cy5-BG

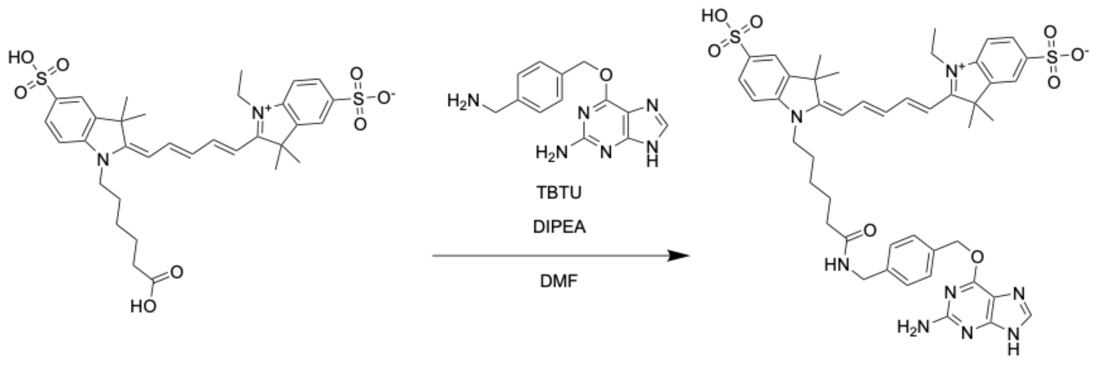


To a vial containing a magnetic stirring bar, sulfo-cyanine 5 carboxylic acid (29.5 mg, 1.0 equiv), 6-((4-(aminomethyl)benzyl)oxy)-9*H*-purin-2-amine (12.1 mg, 1.0 equiv), 2-(1*H*-benzo[*d*][1,2,3]triazol-1-yl)-1,1,3,3-tetramethylisouronium tetrafluoroborate (TBTU) (18.8 mg, 1.3 equiv), *N*,*N*-diisopropylethylamine (DIPEA) (23 μL, 3.0 equiv) and DMF (400 μL) were added. After stirring the mixture for 1 day at 30 °C, 1 N HCl aq. was added. The precipitate was collected by filtration, washed with HCl aq., and dried to afford **Sulfo-Cy5-BG** as a deep-purple-colored solid (18.1 mg, 61% yield). The reagent was used for the SNAP-tag labeling without further purification. $^1$H NMR (500 MHz, DMSO-$d_6$) δ 8.38–8.33 (m, 3H), 7.82 (m, 2H), 7.64 (m, 2H), 7.51 (m, 2H), 7.34–7.25 (m, 4H), 6.53 (dd, *J* = 12.1, 12.1 Hz, 1H), 6.29 (d, *J* = 13.3 Hz, 2H), 5.55 (s, 2H), 4.23 (d, *J* = 4.4 Hz, 2H), 4.13–4.08 (m, 4H), 2.12 (t, *J* = 6.6 Hz, 2H), 1.72–1.68 (m, 14H), 1.57 (t, *J* = 7.2 Hz, 2H), 1.36–1.30 (m, 2H), 1.25 ppm (t, *J* = 6.4 Hz, 3H); $^{13}$C NMR (125 MHz, DMSO-$d_6$) δ 172.75, 172.48, 171.79, 158.31, 154.21, 154.05, 153.10, 152.10, 145.09, 144.82, 141.95, 141.34, 140.49, 140.36, 140.20, 135.87, 133.35, 128.90, 128.70, 127.25, 127.10, 125.95, 125.87, 125.60, 119.84, 119.80, 110.00, 109.94, 109.79, 103.28, 102.99, 68.51, 48.71, 48.65, 45.88, 43.23, 41.56, 39.91, 38.48, 34.76, 26.91, 26.78, 26.50, 25.51, 24.72, 11.95 ppm; HRMS (ESI): *m*/*z* calcd for $C_{46}H_{52}N_8O_8S_2+H^+$: 909.3422 $[M+H]^+$; found: 909.3436.

Note; Peaks of impurities might be overlapped around δ 1.7 ppm in the $^1$H NMR spectral chart.

**Table S1. Primers used in this study.**

| Name | Sequence | |
|---|---|---|
| Nde1_6×HN_SacⅡ_F | GAAGGAGATATACATATGGGTCATAATCATAATCATAATCATAATCATAATCACAACTCCGCGGCTCTTGAAGTCC | Construction of pET47b-6×HN |
| Nde1_6×HN_SacⅡ_R | GGACTTCAAGAGCCGCGGAGTTGTGATTATGATTATGATTATGATTATGATTATGACCCATATGTATATCTCCTTC | Construction of pET47b-6×HN |
| mEGFP_pET47b_Sma1_Sac1_F | CCTCTTTCAGGGACCCGGGATGGTGAGCAAAGGCGAAG | Construction of pET47b-6×HN-NT-mEGFP |
| mEGFP_pET47b_Sma1_Sac1_R | GGCACCAGAGCGAGCTCTTTATACAGTTCATCCATGC | Construction of pET47b-6×HN-NT-mEGFP |
| SNAP_pET47b_Sma1_Sac1_F | CTCTTTCAGGGACCCGGGATGGATAAAGACTGCGAAATGAAAC | Construction of pET47b-6×HN-NT-SNAP |
| SNAP_pET47b_Sma1_Sac1_R | CGTGGCACCAGAGCGAGCTCGCCCAGGCCTGGTTTAC | Construction of pET47b-6×HN-NT-SNAP |
| AtD14_pET47b_Sma1_Sac1_F | CTCTTTCAGGGACCCATGAGTCAACACAACATCTTAGAAG | Construction of pET47b-6×HN-AtD14/AtD14$^{S97A}$-mEGFP |
| AtD14_pET47b_Sma1_Sac1_R | CGTGGCACCAGAGCGAGCTCCCGAGGAAGAGCTCGCCG | Construction of pET47b-6×HN-AtD14/AtD14$^{S97A}$-mEGFP |
| AtD14_pET47b_Xho1_Pac1_F | TTCTGCTGCTCTCGAGATGAGTCAACACAACATCTTAGAAG | Construction of pET47b-6×HN-mEGFP-AtD14 |
| AtD14_pET47b_Xho1_Pac1_R | TAGCAGCCTAGGTTATCACCGAGGAAGAGCTCG | Construction of pET47b-6×HN-mEGFP-AtD14 |
| OmKAI2d3_pET47b_Sma1_Sac1_F | CCTCTTTCAGGGACCCGGGATGTCTACAGTGGGAGCTG | Construction of pET47b-6×HN-OmKAI2d3-(mEGFP or SNAP) |
| OmKAI2d3_pET47b_Sma1_Sac1_R | GGCACCAGAGCGAGCTCAGCAATATCATGTTGGATATG | Construction of pET47b-6×HN-OmKAI2d3-(mEGFP or SNAP) |
| OmKAI2d3_pET47b_Xho1_Pac1_F | CTACTTCTGCTGCTCTCGAGATGTCTACAGTGGGAGCTG | Construction of pET47b-6×HN-(mEGFP or SNAP)-OmKAI2d3 |
| OmKAI2d3_pET47b_Xho1_Pac1_R | TAGCAGCCTAGGTTAATTAATCAAGCAATATCATGTTGGAT | Construction of pET47b-6×HN-(mEGFP or SNAP)-OmKAI2d3 |
| Sac1_GSSG_Xho1_F | CGGTAGCTCAGGAC | Substitution of original linker to GSSG linker |
| Sac1_GSSG_Xho1_R | TCGAGTCCTGAGCTACCGAGCT | Substitution of original linker to GSSG linker |

| | | |
|---|---|---|
| Nco1_3×FLAG_BamH1_F | CCGGCGATGGCCATGGACTACAAGGATCATGATGGGGACTATAAGGATCACGATATTGACTACAAAGATGACGATGACAAGGATCCGAATTCGAGC | Construction of pET26b-3×FLAG |
| Nco1_3×FLAG_BamH1_R | GCTCGAATTCGGATCCTTGTCATCGTCATCTTTGTAGTCAATATCGTGATCCTTATAGTCCCCATCATGATCCTTGTAGTCCATGGCCATCGCCGG | Construction of pET26b-3×FLAG |
| Nco1_Strep Ⅱ -HiBiT_BamH1_F | CCGGCGATGGCCATGGCAAGCTGGAGCCACCCGCAGTTCGAAAAGGGTTCAAGCGGTGTGAGCGGC | Construction of pET26b-Strep Ⅱ-HiBiT |
| Nco1_Strep Ⅱ -HiBiT_BamH1_R | GCTCGAATTCGGATCCTGAGCTACCGCTAATCTTCTTGAACAGCCGCCAGCCGCTCACACCGCT | Construction of pET26b-Strep Ⅱ-HiBiT |
| BamH1_AtSMXL7_Xho1_F (FLAG) | ACGATGACAAGGATCCGATGCCGACACCAGTAACCAC | Construction of pET26b-3×FLAG-AtSMXL7 |
| BamH1_AtSMXL7_Xho1_F (StrepⅡ-HiBiT) | CGGTAGCTCAGGATCCATGCCGACACCAGTAACCACG | Construction of pET26b-Strep Ⅱ-HiBiT--AtSMXL7 |
| BamH1_AtSMXL7_Xho1_R | GGTGGTGGTGCTCGAGGATCACTTCGACTCTCGCCG | Construction of pET26b-(3×FLAG or Strep Ⅱ -HiBiT)-AtSMXL7 |
| BamH1_OmSMAX1_Xho1_F | ACGATGACAAGGATCCGATGAGAGCCGGATCCAGCACTATC | Construction of pET26b-3×FLAG-OmSMAX1 |
| BamH1_OmSMAX1_Xho1_R | TGGTGGTGGTGGTGCTCGAGAACCCTCAACCAATCCTCATTGTCC | Construction of pET26b-3×FLAG-OmSMAX1 |
| BamH1_OmSMAX1-D1M_Xho1_F | ACGATGACAAGGATCCGGCCACCATCGAACAATCATTAACC | Construction of pET26b-3×FLAG-OmSMAX1-D1M |
| BamH1_OmSMAX1-D1M_Xho1_R | TGGTGGTGGTGGTGCTCGAGTTTATATGTACTGGCATCTAAAGTATTAGCAAAC | Construction of pET26b-3×FLAG-OmSMAX1-D1M |
| AtMAX2_pDONR207_F | GGGGACAAGTTTGTACAAAAAAGCAGGCTTAATGGCTTCCACTACTCTCTCCGACC | Construction of pDONR207-AtMAX2 |
| AtMAX2_pDONR207_R | GGGGACCACTTTGTACAAGAAAGCTGGGTATCAGTCAATGATGTTGCGGCTGTTC | Construction of pDONR207-AtMAX2 |
| ASK1_pDONR207_F | GGGGACAAGTTTGTACAAAAAAGCAGGCTTAATGTCTGCGAAGAAGATTGTGTTG | Construction of pDONR207-ASK1 |

| | | |
|---|---|---|
| ASK1_pDONR207_R | GGGGACCACTTTGTACAAGAAAGCTGGGTATCATTCAAAAGCCCATTGGTTC | Construction of pDONR207-ASK1 |
| AtMAX2_pUPD2_F | GCGCCGTCTCGCTCGAGCCATGGCTTCCACTACTCTCTCCGACC | Construction of pUPD2-AtMAX2 |
| AtMAX2_pUPD2_R | GCGCCGTCTCGCTCAAAGCTCAGTCAATGATGTTGCGGC | Construction of pUPD2-AtMAX2 |
